# Continuous Strategy Adaptation and Discrete Switching Driven by Environment and Internal State in Meta-Learning

**DOI:** 10.64898/2026.06.08.729424

**Authors:** Jianning Chen, Masakazu Taira, Kenji Doya

## Abstract

Behavioral strategies can change in response to environmental and internal states, either gradually or abruptly, enabling flexible adaptation. Such strategy regulation is central to meta-learning, the ability to learn to learn. Previous studies analyzed temporal or condition-dependent strategy change using models and theories that assume continuous or discrete changes. Here, we analyze the mice’s behavior in a two-step decision task using four different approaches: stay-switch choice probability analysis; generalized linear mixed model (GLMM) of choice and reaction time (RT) given preceding task events; fitting a reinforcement learning (RL) model with time-varying meta-parameter by a novel multiple-step particle filtering method; and fitting a finite internal state (FIS) model that produces choice and RT depending on discrete state transition. Together, the stay probability and GLMM analyses reveal that learning progress encourages a shift toward a model-based, value-based learning strategy, accompanied by elevated choice perseveration. More uncertain reward settings or changes in them lead to random, exploratory behavior. Meta-parameter dynamics show faster learning, greater involvement of a model-based strategy, higher choice stochasticity, and more rapid development of choice perseveration with less contribution to the final decision as learning progresses. Exploratory behavior in the face of uncertain reward settings or changes in those settings is underpinned by slower forgetting and greater model-based contribution. FIS modeling discovered a trial-level switch between an optimal value-based learning state and a suboptimal self-repeating state. Meta-parameter dynamics reflect continuous strategy changes, while state transitions capture abrupt, discrete strategy switches. At an intermediate timescale, when reward settings change, two processes interact: mice persist in a self-repeating state, leading to attempts at model-based strategy with incomplete adaptation.

## 1 Introduction

Learning and decision-making behaviors do not occur in isolation, nor are they static. Behaviors often exhibit consistency and systematic change, governed by similar computational rules, commonly referred to as behavioral strategies [1]. Two main classes are value-based learning, including model-based reinforcement learning (RL), model-free RL, and the win-stay-lose-shift (WSLS) strategy, and value-free learning, which is primarily instantiated as choice perseveration. Value-based strategies compute value, as described by various RL algorithms [2, 3, 4], whereas value-free learning mainly manifests choice persistence through repetition [4, 5, 6]. Adaptively regulating these learning and decision-making strategies in response to the learning progress [7, 8, 9, 10], environment (e.g., uncertainty [8, 11], task and policy complexity [12, 13, 6, 14, 15]) and internal state (e.g., task engagement, motivation, confidence and fatigue) [16, 17, 18] appears in distinct timescales, mainly in a gradually continuous [19, 2, 7, 15] or abruptly discrete [16, 18, 10, 20, 21, 22] manner. Learning to select appropriate strategies for learning and adapting to current settings is an example of meta-learning, which refers to the case where experience assists fast learning [23, 24, 25, 11]. Several key examples include shifts in model-based and model-free control [8, 26, 12, 13], exploration-exploitation tradeoff [27, 28, 24, 21], and the balance between reward-seeking and choice perseveration tendencies [6, 14, 5].

Multiple factors shape the strategy regulation. Certain environmental factors modulate behavioral strategies directly, such as uncertainty [8, 11] and resource availability [29]. Likewise, some aspects of internal states influence behavior strategies independent of the environment, including fatigue [30, 31] and satiety [32]. Occasionally, the extent and manner in which environmental changes influence strategy selection depend on how the internal state responds to those changes. For example, studies observed that increasing task difficulty may lead to the use of sophisticated strategies or to aversion to it, potentially depending on whether decision-makers can and are willing to invest effort [33, 34, 13, 35].

Various computational approaches to investigate the strategy changes have been developed, mainly assuming either continuous or discrete changes. In RL-based frameworks, strategy changes are represented by continuous adjustments to meta-parameters [7, 19, 2, 15]. Another class of methods utilizes the hidden Markov process and views the strategy changes as a discrete state transition [16, 18]. Two lines of research merge, motivating an investigation of whether and how two processes coexist, and whether they interact in shaping behavior strategy regulation.

In this paper, we conduct a mouse two-step task [26, 3, 36] and apply four approaches with improvements to investigate the changes in behavioral strategies at multiple timescales. The stay-switch choice probability provides an overview description of choice patterns. We refine the logistic regression model [26, 37, 3] to separate the effects of task events on choice and reaction time. We also developed a multi-step particle-filtering approach for dynamic RL modeling. The estimated time-varying meta-parameter reveals strategy adaptation at the long timescale in a continuous manner. Rather than relying solely on the choice [18, 10, 16], we propose a novel hidden Markov model with a finite internal states model (FIS) that jointly models behavior and response time to identify internal states that reflect how strategies switch discretely in the trial resolution. Together, applying these methods, we studied how behavioral strategy regulation is modulated by environment, internal state, and their interaction, integrating continuous and discrete perspectives on strategy change. These empirical and methodological results would contribute to an understanding of behavioral adaptation across different timescales.

## 2 Result

### 2.1 Task Description and analysis tool validation

10 Tph2-tTA mice performed the two-step task [36] (figure.1.a). Mice were first asked to choose between the left and right options, followed by the presentation of the second-step state (up or down), depending on the first-step choice and the predetermined transition probability (either common 80% or rare 20%). The up and down states might lead to a reward with a probability that varies across the three types: good/bad, bad/good, and neutral/neutral, corresponding to up, down, and neutral blocks. The correct decision is the option that commonly leads to the state with the higher reward probability. After training, mice were able to optimize their choices to the reward settings(fig.1.b). Mice were trained by stages, and we included the last two training stages (I & II) and the testing stage (III) (4.3) in the main analysis. In these stages, mice only move to the next if they can perform a sufficient number of blocks per session for constitutive days (4.3). Mice usually perform 2-3 days in stage I, 3-5 days for stage II, and 12 days for stage III.

**Figure 1.**
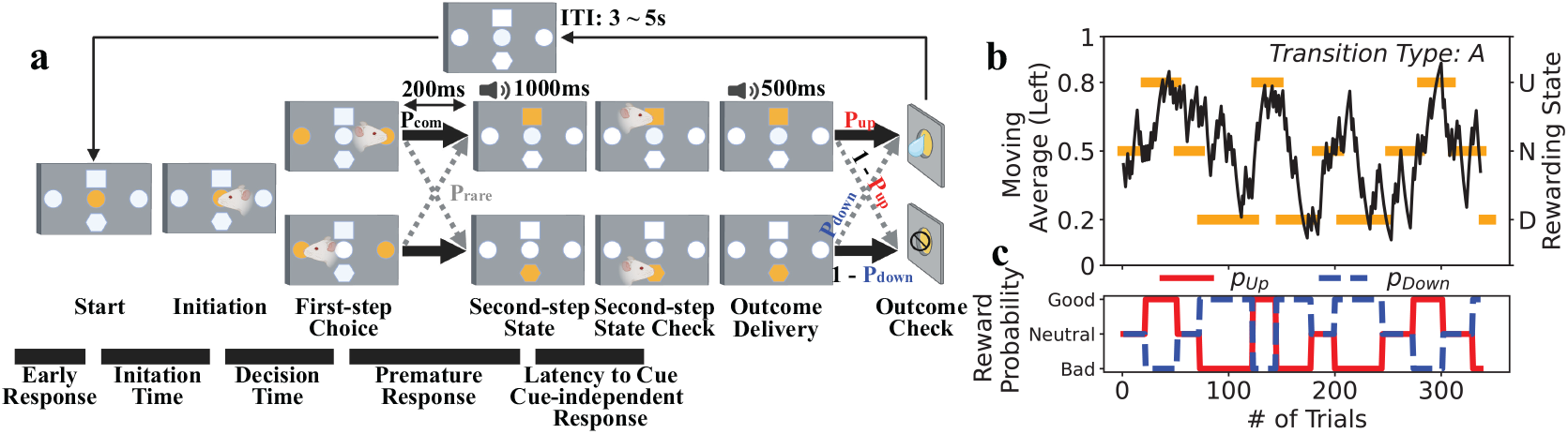
Task structure and example behavior. **a**, two-step task and response time measurements (see 2.4). After the inter-trial interval (ITI) lasting 3 *∼*5 seconds, the next trial starts, indicated by the illumination of the center port. After initiation by nose-poking the center port, a first-step choice between left and right is presented by the illuminated ports in free-choice trials. In forced-choice trials (not shown here), only one port (option) is illuminated (available). Following a 200ms delay after the choice, the up or down second-step state is reached with either common or rare probability, as indicated by the illumination of either the up or down port, accompanied by a 1000ms state cue. After the second-step state check by nose-poking the illuminated port, outcomes are delivered with a 500 outcome cue, at different probabilities. Several response time measurements are extracted: 1) early response before the trial starts; 2) initiation time between the trial starts and the first-step choice; 3) premature response before the second-step state is presented; 4) latency to cue spent on second-step state check; and 5) cue-independent response responds to the unvisited state. **b**, Example behavior from one session with the A-type transition matrix. The exponential moving average of the animal’s left choice (black line) tracks the current rewarding state (orange bar). **c**, the reward probability for up and down states changes anti-correlated. In the non-neutral block, the block changes once the accuracy reaches 0.75, whereas block changes occur after 20-30 trials in the neutral block (4.3).

### 2.2 Learning progress and reward settings modulate the stay probability patterns

We first apply the stay probability analysis to investigate the behavior, which examines the frequency of choosing the same action as the last trial after four trial types with different transition types and outcomes (i.e., CR, common/reward; RR, rare/reward trials; CN, common/no reward; and RN, rare/no reward).

To examine whether learning progress drives strategy adaptation, we focus on mice’s behavior in the last three stages. The overall stay probability increased (stage I: 58.1969% ±4.6812%; stage II: 63.2090%±3.7870%; stageIII: 67.5948% ±2.3721%) as the stage proceeded (fig.2.a). Experiencing more trials strengthens choice perseveration at stage III, possibly due to the habituation via stimulus-response pairing (fig.2.a). Starting from stage II, mice learned to stay often after CR events, suggesting that value-based learning took place (fig.2.b). However, such a tendency is modulated by the transition type. Mice switch the most frequently after RR event, exhibiting the “U-shape” stay probability distribution (fig. 2.b). Despite other notable differences, this pattern is similar to that generated by model-based strategies (Supplementary fig.1.a). The change in stay probability over the training stage suggests that the use of a model-based strategy may change with learning progress, yet whether other changes in strategies co-occur remains unclear. In stage III, when performance is stabilized, mice also seem to continuously adjust their strategies in response to the reward settings and their changes. The “U-shape” stay probability distribution, which might imply the involvement of model-based strategies, is flattened in the neutral block (13592 trials) compared with other types (42577 trials) (fig. 2. c) or 10 trials after the block type changes (13592 trials) compared with the rest of trials ((39944 trials)) (fig. 2. d). It suggests that mice may lean toward a simpler, model-free strategy when reward settings change or remain neutral.

**Figure 2.**
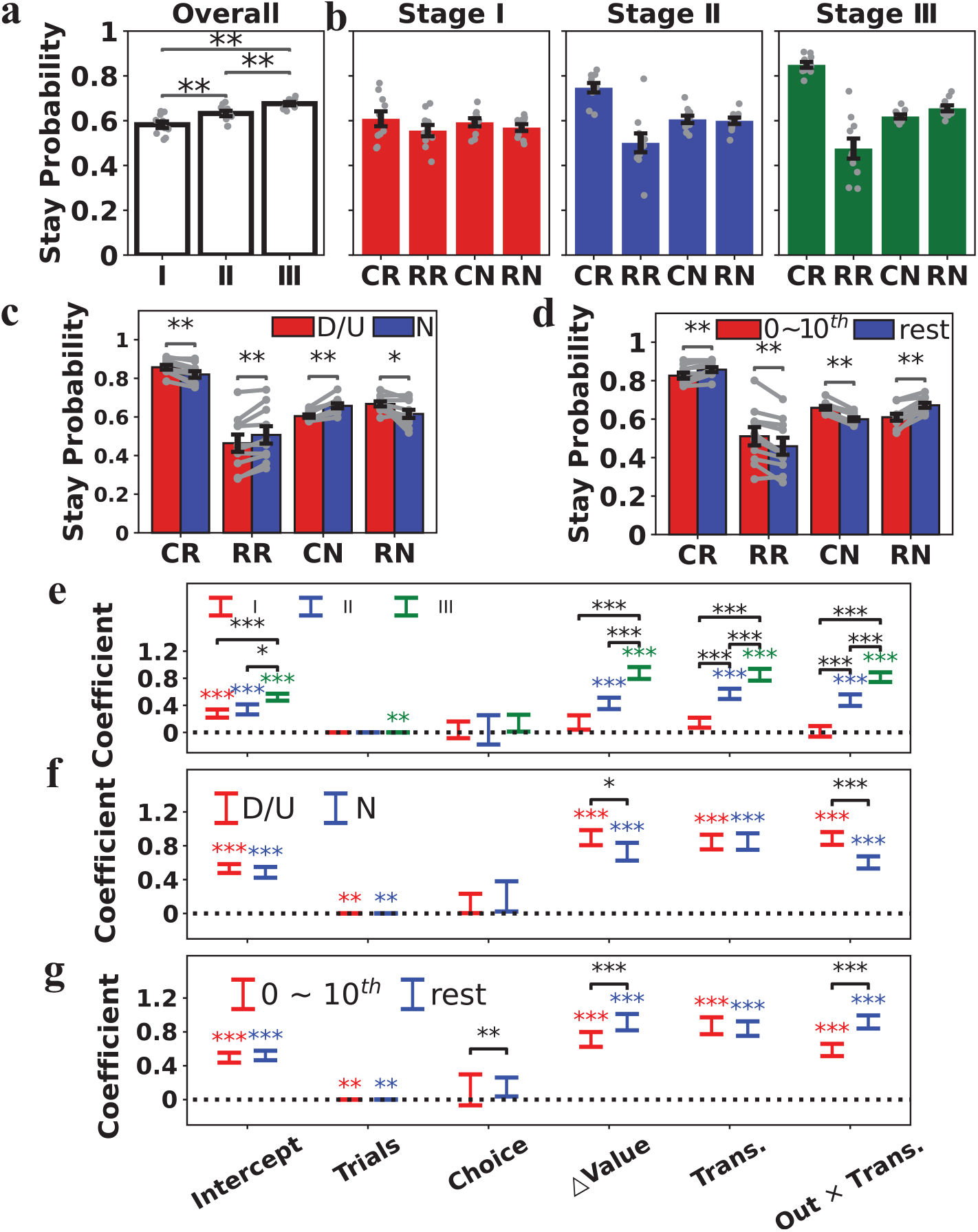
Behavioral analysis result. **a**, the overall stay probability in three learning stages. The error bars show the mean and S.E.M, which applies to all following figures, unless specified otherwise. Statistics are obtained from the Wilcoxon test. **b**, stay probability analysis in three stages. The first three panels show the stay probability after four trial types (i.e., CR, common/reward; RR, rare/reward; CN, common/no reward; and RN, rare/no reward). **c**, stay probability in non-neutral blocks, up (U) and down (D) blocks, and neutral blocks (N). **d**, stay probability in the first 10 trials after the change and the rest of the trials. **e**, coefficient results from the GLMM for each stage and between stages. Statistical significance between stages is obtained by the interaction of stages and the respective regressor (4.4.2). **f**, GLMM result for block type comparison. **g**, GLMM result for the comparison of one behavior in the first 10 trials after the change and the rest of the trials.

### 2.3 GLMM analysis reveals the changes in strategies underlying the behavioral changes

To further investigate the underlying strategy, we apply logistic regression [26, 3, 36], which can differentiate the effects of previous task events and choices on the current decision. We refine and validate the regression model structure (see the model specification in 4.4.5 and the validation results in the Supplementary.1). Besides the left or right choice (Choice), whether common or rare transition experienced (Trans.) and the reward-transition interaction (Out× Trans.) showing whether model-free or model-based system demonstrate, the final structure introduces the number of trials experienced so far (Trials) to account for the possibility that choice perseveration might be exacerbated by habituation from repeated presentations of identical content or as time passes [38, 39, 4]. It also includes the inferred value difference (∆Value), a cognitively plausible approximation of the reward history. It is the difference in reward probabilities between the chosen and unchosen options estimated from the temporally discounted outcomes history (4.4.2).

Consistent with the increasing overall stay probability, experiencing more trials leads to more stay decisions at stage III, suggesting that strengthened choice perseveration is possibly due to habituation via stimulus-response pairing (fig.2.e). Animals learned to repeat the choice associated with the better reward history, shown by an increasing coefficient of inferred value difference (stage I: *β*_∆*Value*_ = 0.1465 ± 0.1057, *p* = 0.1661; stage II: *β*_∆*V alue*_ = 0.4288 ± 0.0843, *p* < .0001; stageIII: *β*_∆*Value*_ = 0.8772 ± 0.0876, *p* < .0001) as the stage proceeded (fig. 2.e). Consistent with the emergence of the “U-shape” stay probability distribution, transition modulates mice’s choice in pursuing the reward, which developed over stages, leading to the growing coefficient in the Out × Trans (stage I: *β*_*Out* × *Trans*._ = 0.0158± 0.00776, *p* = 0.8383; stage II: *β*_*Out* × *Trans*._ = 0.4765 ±0.0849, *p* < .0001; stageIII: *β*_*Out* × *Trans*._ = 0.8162 ± 0.0712, *p* < .0001; fig.2.e). The model-based system gradually dominates the mice’s behavior as learning proceeds. Beyond what the stay probability suggested, a significant coefficient of transition type in stage II and III (fig.2.e) resembles the fictive learning [40]. In brief, the fictive learning updates the value of the unvisited state by generalizing the reward prediction error scaled by the estimated correlation between visited and unvisited states.

Compared with non-neutral blocks, in neutral blocks, mice exhibit more exploratory behavior, as indicated by a decreasing coefficient for ∆Value (*β*_∆*Value*:*blocktype*_ = −0.1711± 0.0760, *p* = 0.0244), with less model-based planning, suggested by the lower Out × Trans (*β*_*Out* × *Tran*:*blocktype*_ = −0.2854 ± 0.0567, *p* < .0001, fig.2.f). The same changes were found in the first 10 trials after the block changes (16225 trials), comparing to the rest of trials (39944 trials)(*β*_∆*Value*:*Laterphase*_ = 0.2627 ± 0.0768, *p* = 0.0006. *β*_*Out* × *Tran*:*Laterphase*_ = 0.3046 ±0.0484, *p* < .0001, fig.2.d & g).

To better understand mice’s behavior in neutral blocks or right after a block change, when information about current reward settings is limited, we examine the hypothesis that mice’s behavior in these conditions is partially guided by information from the preceding blocks. That is, whether mice tended to choose the options that were the correct choice in the last non-neutral block. Overall, mice, in neutral blocks, tended to choose these more than chance level (58.8317 ±4.2535%, *W* = 54, *p* = 0.0020, Wilcoxon test), which is the opposite of that in non-neutral blocks (37.4252± 2.6784%) due to learning. Such dependence quickly disappears after the first 10 trials in a non-neutral block (from 55.6563 ±6.1905% to 32.0058 ± 3.8057%), suggesting that adaptation was in progress. By contrast, in the neutral block, this dependence was slightly mitigated yet remains above-chance level (from 55.6563±6.1905% to 54.7516±4.7649%), implying incomplete adaptation. In general, these observations suggest that mice tend to use outdated information to guide their decision when information is lacking.

Combining both stay probability and GLMM analysis, over the course of learning, the mice continuously adjust their behavioral strategy from seemingly random to choice-preserved, exploitative, and model-based strategies with fictive learning. When facing neutral reward settings or changes in them, more exploratory behavior with a less model-based signature emerges.

### 2.4 Response time patterns reflect the internal state

Aspects of the internal state, such as fatigue, motivation, satiety, and task engagement, may modulate the use of learning strategies. These factors are usually hidden, which we characterize by the reaction time (RT). A typical trial consists of trial initiation, first-step choice, and second-step state check, yielding rich RT-related measures that reflect distinct cognitive activities and aspects of internal state (fig.1.a). We derive measures and then examine how those measures are related to specific behavioral aspects and task events (4.4.2), to reveal the related cognitive activities and internal states. The selected measures are *early response, initiation time, decision time*, and *latency to cue* (4.4.1). To better separate the effect of learning from the current analysis, the main GLMM analysis were restricted to Stage III, when performance was stable as per training instructions (4.3).

The animal occasionally attempts to initiate the trial during the ITI, termed the early response. More early responses occurred as the stage progressed (I:27.0129% ± 13.2205%; II: 34.3415% ± 20.2937%; III: 44.0640% ± 17.4239%), implying that it might represent the elevation of confidence and motivation through learning rather than ignorance of the instruction. In stage III, more early responses appear in the later phase in a block (first 10 trials: 41.4653% ± 16.6780%; rest: 45.1190% ± 17.7520%) and after successfully obtaining more rewards in the past (fig.3), consistent with its role in reflecting confidence. Besides, earlier responses are observed as the reward history improves, which decreases over time (fig.3), suggesting it might signal high motivation and low fatigue. An early response might also indicate high impulsiveness and choice readiness, as it is likely to occur during longer ITIs and before making the same choice (stay: 46.2121% ± 18.2259%; switch: 40.4081% ± 16.1367%) (fig.3).

**Figure 3.**
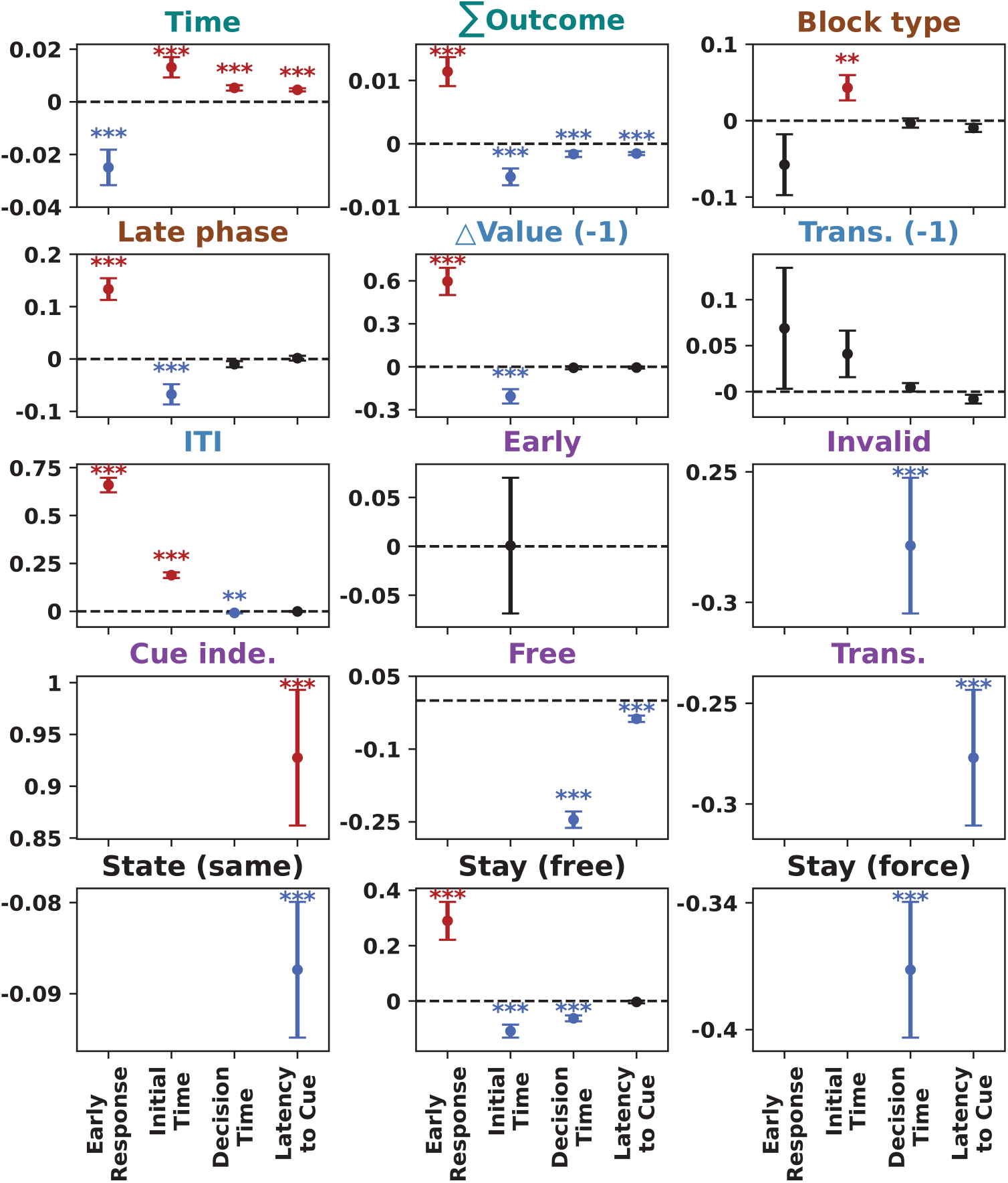
Reaction time analysis and measure selection. GLMM models for RT-related measures, using a logit and log link functions for early response and other measures, respectively. Predictors are grouped by the functional meaning (4.4.2). Long-term effects (teal): time (seconds elapsed) and ∑ Outcome (number of rewards obtained). Block-wise effects (saddle brown): neutral (if in a neutral block) and late phase (if 10 trials have passed since the block changes). Last-trial effect (steel blue): ∆Value (-1) (inferred value difference in the last trial), trans. (-1) (transition type in last trial), and ITI (the inter-trial interval). Current-trial effect (dark violet): early (if the early response is made), invalid (if the invalid choice has been made in the force-choice trial), cue. indep. (if the cue-independent response has been made), free (if it is the free-choice trial), and tran. (current transition type). Effects of the contrast between the last and current trial (black): state (if the same state was presented both), stay (free) (if stay in the free-choice trial), and stay (force) (if stay in the force-choice trial). Each sub-panel shows the GLMM coefficient for one regressor. Significant coefficients are colored red if positive, or blue if negative, while coefficients without significance are black.

The initiation time for proper or early responses captures features similar to those of the early response. Mice quickly initiated the trial in the later stage (I:2.2724 ± 0.9049 s; II: 1.8545 ± 0.5956 s; III: 1.4922 ± 0.1868 s, all reaction times are reported as mean ± SD of subject-wise median values), and later phase of the block (first 10 trials: 1.5529 ± 0.2096 s; rest: 1.4728 ± 0.1872 s), associated with more obtained reward, a higher inferred reward difference, and the stay choice (stay: 1.4553 ± 0.1750 s; switch: 1.5621 ± 0.2189 s), but showed longer initiation time in neutral block(neutral: 1.5379 ± 0.1727 s; non-neutral: 1.4798 ± 0.1938 s) or after spending long time in the session (fig.3), suggesting that a short initiation time might reflect high confidence, motivation and choice readiness and low fatigue. However, unlike the early response, longer ITIs were often followed by longer initiation times (fig.3), suggesting disengagement from the task.

Decision time might reflect choice confidence, readiness, and fatigue. Learning progress shortens the decision time in both free (I:1.0281 ± 0.1444 s; II: 0.7916 ± 0.1622 s; III: 0.5906 ± 0.0681 s) and forced-choice trials (I:1.5342 ± 0.5105 s; II: 1.1866 ± 0.5545 s; III: 0.8756 ± 0.1463 s) (Fig.3), possibly by promoting the confidence. Similarly, higher confidence, encouraged by earning more rewards, leads to shorter decision times (Fig.3). Additionally, fatigue from time spent in the experiment slows decision time (Fig.3). No significant coefficients are found for the regressors related to the motivation for the chosen option (∆Value) and task difficulty (Block type and Later phase).

Latency to cue is the time spent in the second-step state check, with two types of erroneous responses: premature and cue-independent. The animal often prematurely intends to start a trial, before it is ready, during the delay, and the tone (I:77.7574% ± 2.3350%; II: 89.1658% ± 3.1622%; III: 95.1444% ± 1.7174%), suggesting eagerness for reward. Animals also occasionally check the state not presented (I:19.9165% ± 3.1134%; II: 10.8556% ± 2.2459%; III: 4.5952% ± 1.3267%). The cue-independent response occurred mainly after an unexpected transition (i.e., stay the same choice, but arrived at a different state) (I:47.2335% ± 10.6853%; II: 36.9853% ± 11.5314%; III: 19.6832% ± 8.8542%). It suggests that the cue-independent response is likely to be a reflex. Therefore, latency to cue is defined as the time required to confirm the second-step state for both valid and premature responses, but not for the cue-independent response. The latency to the cue is only marginally modulated by long-term confidence and fatigue, but highly depends on whether the cue is predictable (common: 0.5161 ± 0.0053 s; rare:0.7280 ± 0.0844 s) (Fig.3).

### 2.5 The dynamics of meta-parameter in RL models underlying the observed behavioral changes

To uncover the computational principle underlying these observed behavioral changes due to learning progress and reward settings, we conducted RL-based modeling to estimate the meta-parameter dynamics, assuming the continuous strategy adaptation. Conventional modeling estimates only static meta-parameter distributions. A common workaround is to fit each session or block independently to identify the best-fitting model and parameter values. However, by fitting 61 models with constant parameters, we found different sessions often favor different models (Supplementary.2), and meta-parameter estimates are inherently model-specific, rendering cross-model comparisons of meta-parameters incomparable. These observations highlight the importance of the dynamic modeling approach.

The model for dynamical investigation includes computational rules and meta-parameters that improve fit in fixed-parameter modeling. Consistent with previous two-step task literature, we use the hybrid model between model-based (MB) and model-free (MF) systems, with *ϵ* representing the weight of the MF system relative to the MB system. In the MF system, we include the eligibility trace between action and state (i.e., *λ*). Additionally, fictive learning (FL), forgetting (forget), and choice perseveration (Perse) are included. The forgetting rule (forget) assumes that, for the action or state that was not experienced in the last trial, its cached value decreases by a certain forgetting rate, *α*_0_. Choice perseveration (Perse) learns how often an action has been chosen by learning rate *α*_*CK*_, with contribution to the final behavior scaling by *ϵ*_*CK*_. Fictive learning (FL) assumes that the agent can generalize the actual, experienced prediction error to the unchosen action or state that was not presented, by scaling the prediction error with event correlation *η*.

We extended the particle-filtering algorithm of [19] to capture the temporal dynamics of meta-parameters. The original algorithm maintains a cloud of particles, each representing a combination of meta-parameters and hidden variables. At each trial, every particle is resampled based on its weight, which reflects the discrepancy between its prediction and the observation. Yet, multiple parameter combinations can yield similar single behavioral output, whereas the genuine probability is unknown, leading to identifiability issues and overfitting.

To address these issues, we propose a forward particle filtering method with an adaptive forward step that evaluates the choice probability based on local consistency. The choice probability and the meta-parameter in local adjacent trials are approximately stationary. The meta-parameters are weighted by their predictions in the next several trials (i.e., forwarding the current meta-parameter to predict future trials). The temporally discounted sum of weights guides the particle resampling. The observations from adjacent trials constrain meta-parameters by informing the choice probability. The forward step also prevents overfitting to the current trial. In simulations with time-varying meta-parameters, incorporating the forward step improves recovery of the meta-parameter trajectory (change in root-mean-square error: −0.1490 ± 0.0872, Supplementary.3). The length of the forward step and the temporal discounting factor are adaptive to the local continuousness 4.4.6.

Behavioral analysis indicates that learning progress might encourage choice perseveration, exploitation, and model-based planning. The strengthened choice perseveration is consistent with the rapid development of choice perseveration with a high learning rate (*α*_*ck*_, stage I: 0.1424 ± 0.0949, stage II: 0.1750 ± 0.1107, stage III: 0.2258 ± 0.1441, fig.4.b). Rapid knowledge updating might assists the exploitation, resulting from the increasing learning rate (*α*, stage I: 0.2886 ± 0.1872, stage II: 0.3914 ± 0.2277, stage III: 0.4123 ± 0.1875) and forgetting rate (*α*_0_, stage I: 0.4519 ± 0.1466, stage II: 0.5249 ± 0.1687, stage III: 0.5386 ± 0.2029, fig.4.b), which got further developed by the decreasing interference of choice perseveration (*ϵ*_*ck*_, stage I: 1.6745 ± 0.9373, stage II: 1.2200 ± 0.5675, stage III: 1.2624 ± 0.8169) (fig.4.b). The weakened model-free weight (*ϵ*, stage I: 0.4667 ± 0.1444, stage II: 0.4592 ± 0.1703, stage III: 0.4013 ± 0.2354, fig.4.b) is also coherent with the dominating role of the model-based strategy in the later learning stage suggested by stay probability and GLMM analysis. As learning process, the event correlation for fictive learning stays negative and continues decreasing (*η*, stage I: −0.0447 ± 0.2354, stage II: −0.0577 ± 0.1772, stage III: −0.1661 ± 0.2782, fig.4.b), implying that the animal learns the anticorrelation along with the learning of the task and the use of model. Additionally, the declining inverse temperature also suggests the possible exploration that develops as learning proceeds (*β*, stage I: 2.3688 ± 0.5718, stage II: 2.3237 ± 0.5260, stage III: 2.0085 ± 0.8175, fig.4.b), which is not shown in previous behavioral analysis.

**Figure 4.**
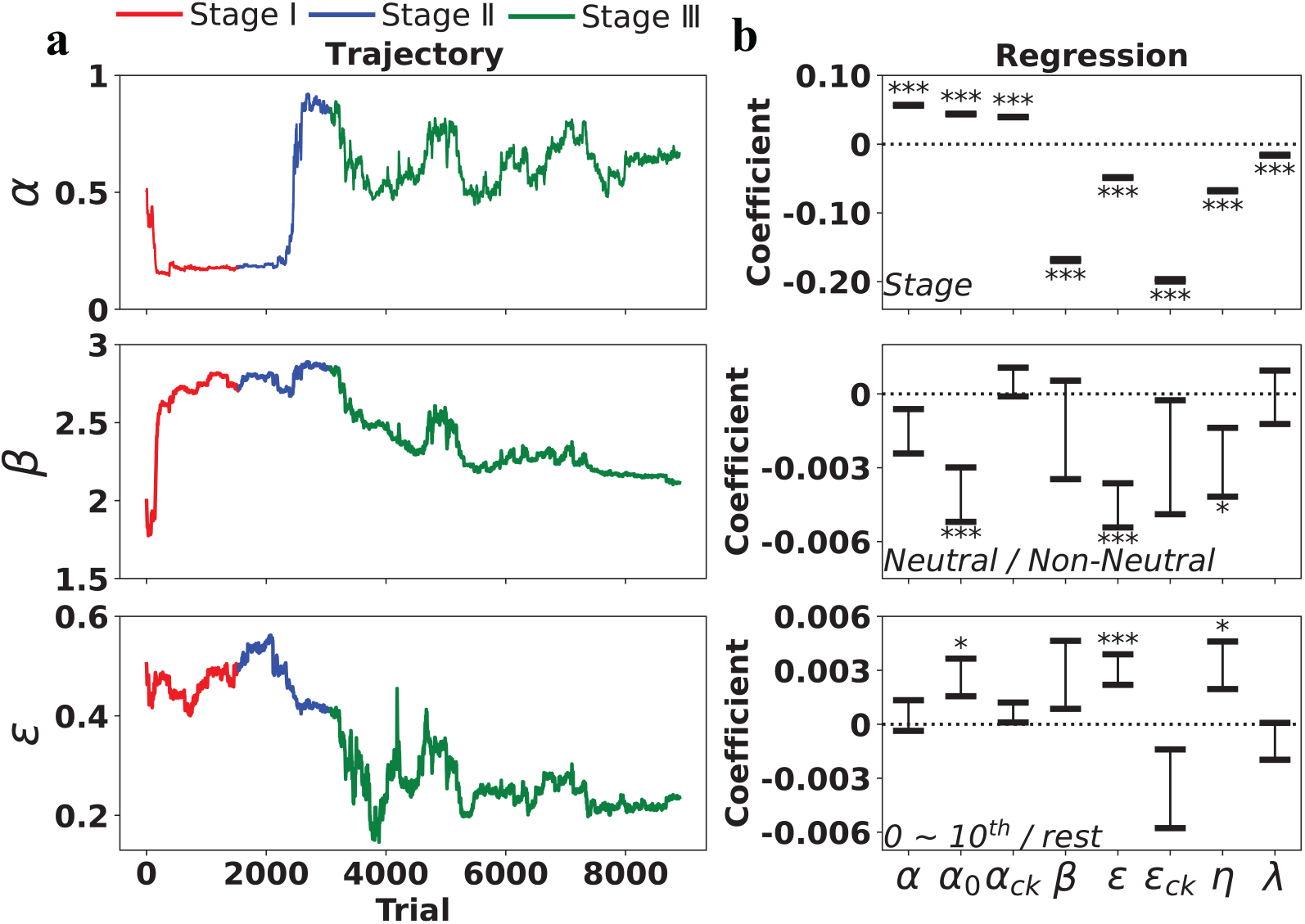
Meta-parameter analysis. **a**, example filtered meta-parameters in three learning stages, three important meta-parameters are selected for illustration. **b**, the GLMM results of meta-parameter predicted by learning stage (up), block type (middle), and change of block type (down). Each error bar shows the GLMM coefficient (estimate and S.D) for each parameter.

Mice show exploratory, model-free behavior in neutral blocks and the first 10 trials after the block change. In neutral block, this behavior appears along with the low forgetting rate (*α*_0_, non-neutral block: 0.5397 ± 0.2019, and neutral block: 0.5352 ± 0.2060), a lower model-free weight (*ϵ*, non-neutral block: 0.4043 ± 0.2355, and neutral block: 0.3921 ± 0.2351), and a low event correlation (*η*, non-neutral block: −0.1646 ± 0.2750, and neutral block: −0.1708 ± 0.2880) (fig.4.b). Similarly, immediately after the block change, we found a similar low forgetting rate (*α*_0_, first 10 trials: 0.5337 ± 0.2036, and rest: 0.5406 ± 0.2026), a lower model-free weight (*ϵ*, first 10 trials: 0.3932 ± 0.2332, and rest: 0.4046 ± 0.2362), and a lower event correlation (*η*, first 10 trials: −0.1659 ± 0.2834, and rest: −0.1662 ± 0.2761) (fig.4.b). The RL modeling results appear to conflict with the stay probability and GLMM analyses.

### 2.6 Reward, correctness, and time elapsed modulate the discrete transition between state and corresponding strategies

We then examine how the internal state might affect the strategy regulation. Since these factors are likely to change abruptly at short timescales, we adopt a computational approach that assumes discreteness. We propose a novel finite internal state (FIS) model based on response time (RT) related measures to jointly characterize the dynamics of internal

For interpretability, four states are characterized as neutral, optimal, choice-repeating, and alerted states based on their characteristics (Supplementary.5). Briefly, neutral states show a chance-level of stay probability and an average-level of RT means. An altered state is similar to a neutral one but has faster latency to the cue. They also show similar chance-level accuracy, proportion of correct responses (Neutral: 50.9444%±13.8196%, altered: 56.8805%±7.9575%). The optimal state achieves the highest performance (70.8143% ± 4.0460%) with fast initiation but longer decision time. The choice-repeating state shows the highest stay probability with short decision, while the accuracy is slightly above chance level (63.2089% ± 2.6635%).

The behavioral patterns in optimal and choice-repeating states primarily reflect optimal value-based and suboptimal self-repeating strategies, whereas neutral and alerted states likely result from varying levels of engagement. Animals spend more trials in optimal (23.9759% ± 10.9176%) and choice-repeating states (73.5819% ± 10.2198%) than the neutral (0.2534% ± 0.2160%) and altered states (2.1888% ± 1.4991%), suggesting that the behavioral changes are primarily due to a shift in behavioral strategy, which then becomes the focus of the analysis (fig.5.b). State transition occurred in 28.6362% ± 8.6281% trials per session on average with relatively short dwell time (3.5304 ± 5.2799), yet this frequency decreased as experiencing more sessions (*ρ* = −0.284, *p* < 0.001, Spearman’s correlation) or in the second half of the session (30.4977% ± 7.1757% and 27.8301% ± 7.1464%, *z* = 3.000, *p* = 0.0098). Learning strategy persists to sustain consistent behavior over short timescales, but is modulated by learning and adaptation.

**Figure 5.**
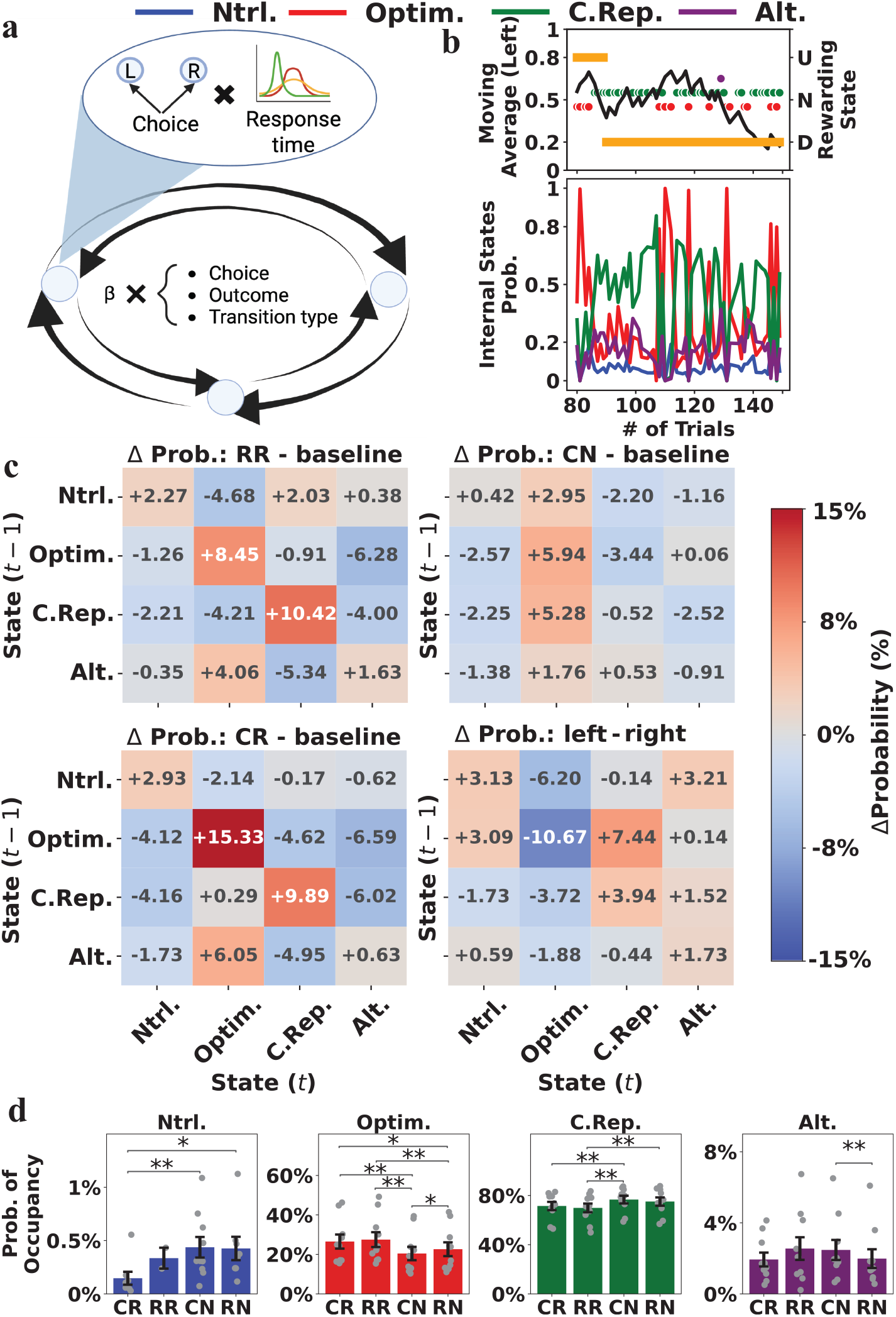
Finite state model and state transition analysis. **a**, the finite state (FIS) model. Each state emits the choice and the response time measures. The transition probability is determined by the choice, transition type, and outcome in the last trial. **b**, the example behavior, and internal state dynamics. The upper panel shows the moving average of the left choice (black line), the rewarding state (yellow bar), and the inferred internal state (colored point). The down panel shows the estimated probability of the internal state. **c**, transition probability changes were reconstructed from the modeling result. The first three panels show deviations following RR, CN, and CR trials relative to RN (baseline), and the last panel shows the change following left versus right choices. **d**, the occupancy rate of the internal state, from the state and the switch in learning strategy. The FIS model (4.4.3) (fig.5.a) assumes that animals transition between states depending on the choice (left or right), the transition type, and the outcome of the last trial. Each state has its own emission probabilities for four RT measures and choices. The time-dependent input (e.g., number of trials, real timestamp) and outcome-transition type interaction are omitted deliberately to examine whether hidden state transitions exhibit intrinsic dynamics in response to these variables without prior assumptions. The model was fitted hierarchically, with subject-level random effects. The model with four hidden states achieves the lowest BIC score (*BIC* Score = 9.7785 × 10^7^) and will be the main focus.

A reward signal may be a general reinforcer for the next choice, the learning strategy, and task engagement. Reward acquisition encourages staying in the same state, as reflected by positive diagonal values following RR and CR events, compared with the RN baseline (fig.5.c). Reward discourages the transitions to the neutral state, leading to its lowest occupancy after CR events (0.1466% ± 0.1716%, fig.5.d). As the inferred value difference increased, the estimated probabilities of the neutral (*ρ* = −0.0165, *p* < .0001, Spearman’s correlation) and alerted states (*ρ* = −0.1748, *p* < .0001, Spearman’s correlation) decreased, suggesting an enhancement of engagement by reward.

The reward also modulated the switching between suboptimal self-repeating and optimal value-based strategies. Following reward delivery, occupancy increased in the optimal state (CR: 26.5001% ± 11.3176%, CN: 20.4362% ± 10.6752%, *w* = 0, *p* = 0.0020, Wilcoxon; RR: 27.4865% ± 11.9109%, RN: 22.6173% ± 11.0702%, *w* = 0, *p* = 0.0020, Wilcoxon) but decreased in the choice-repeating state (CR: 71.4483% ± 10.6660%, CN: 76.6556% ± 9.9213%, *w* = 0, *p* = 0.0020, Wilcoxon; RR: 69.8964% ± 11.2202%, RN: 75.0582% ± 10.6697%*w* = 0, *p* = 0.0020, Wilcoxon.). Consistently, the positive and larger inferred value differences ∆Value occur likely with the optimal state (*ρ* = 0.1033, *p* < .001, Spearman’s correlation) but unlikely with the choice-repeating state (*ρ* = −0.0195, *p* < .0001, Spearman’s correlation).

Similarly, a correct choice in the last trial is followed by a similar shift toward the optimal state (correct: 26.4108% ± 10.8200%, erroneous: 21.9161% ± 11.4779%, *w* = 0, *p* = 0.0020, Wilcoxon) from the choice-repeating state (correct: 71.4090% ± 10.2035%, erroneous: 75.2807% ± 10.6581%, *w* = 0, *p* = 0.0020, Wilcoxon). In contrast, time elapsed within a session possesses the opposite effect. The probability of the choice-repeating state gradually increased (*ρ* = 0.0461, *p* < 0.0001, Spearman’s correlation), whereas that of the optimal state decreased (*ρ* = −0.0347, *p* < 0.0001, Spearman’s correlation), reflecting the strategies deviating from the optimal, possibly due to fatigue and loss of motivation.

### 2.7 Linking the discrete state transition with adaptive meta-parameter adjustment

The RL model reveals how strategy regulation on long timescales (e.g., between sessions) is represented by the changes in the meta-parameter. In parallel, the FIS model shows that switching between optimal and choice-repeating is the primary driver of the behavioral dynamics observed at short timescales. Both perspectives are driven by the same set of observations, and we examine whether they are complementary. We calculate the difference in inferred state probability between the optimal and choice-repeating state and examine how the meta-parameter dynamics might be related to state transition.

The transition from choice-repeating to optimal state marks a shift in control from choice perseveration to value-based and model-based learning. Optimal states are often accompanied by a higher learning rate, exploitation driven by a low inverse temperature, and an increasing weight for the model-based strategy (Fig. 6.a). Meanwhile, the choice-perseveration system is suppressed to ensure the dominant role of value-based learning (fig. 6.a). This result links discrete strategy switching to adaptive adjustment of the meta-parameter, suggesting that the strategy change affects both choice patterns and reaction times.

**Figure 6.**
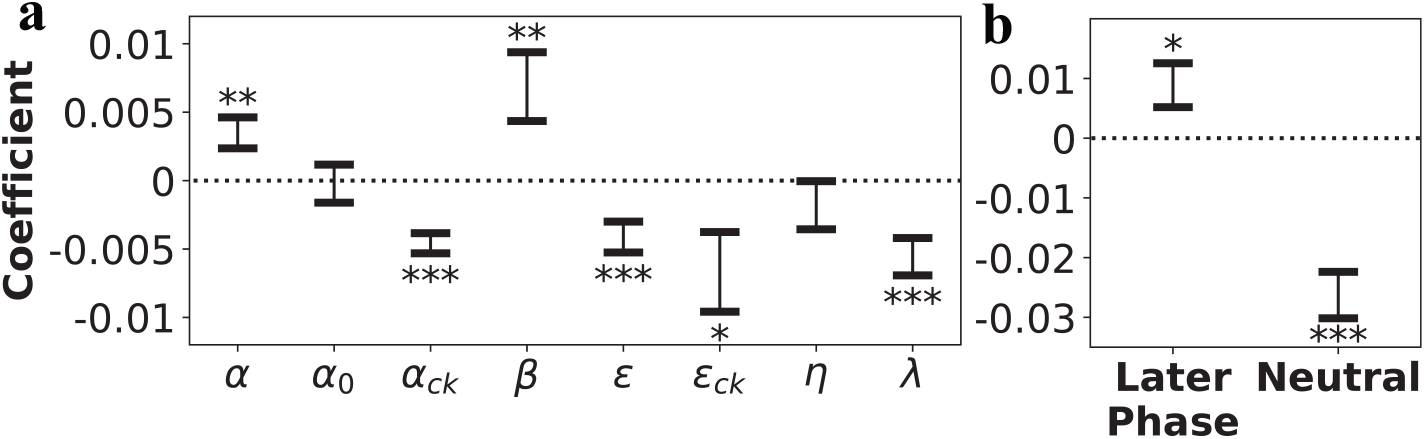
The integrative analysis of meta-parameter dynamics and internal state. **a**, the GLMM result of filtered meta-parameter, predicted by the difference of state probability between optimal and choice-repeating state. **b**, the GLMM result of the difference in state probability between optimal and choice-repeating state, predicted by whether the block changed in the previous ten trials (later phase) and whether it was the neutral block (neutral).

We further ask, for the behavioral strategy change that occurs at the intermediate timescale, between blocks or around block-type changes, whether two processes might jointly shape the behavior. The optimal state likely appears later in a block (*β* = 0.0071 ± 0.0019, *p* = 0.0001) and in a non-neutral block (*β* = −0.0137 ± 0.0020, *p* < .0001) (fig.6.b). Together, after a block change or in the neutral block, when information about reward settings is sparse, mice tended to be in a choice-repeating state and to apply the model-based strategy with slower forgetting.

## 3 Discussion

This study provides evidence for understanding how behavioral strategies change by multiple approaches to elucidate the computational mechanisms. The stay probability analysis, GLMM analysis, and RL modeling consistently suggest that learning progress drives the strategy toward exploitative, model-based fictive learning, with rapid development of choice perseveration and reduced exploration. In trial resolution, FIS and RL modeling jointly reveal the switch between an optimal model-based, exploitative state with fast learning and a suboptimal self-repeating state with strong choice perseveration, modulated by motivation, confidence, satiety, and fatigue. Yet, different approaches yield conflicting results when examining behavior under the neutral and changed reward settings, which warrants further discussion. This study also introduces methodological developments of broad relevance to the neuroscience community. Adaptive particle filtering effectively reveals the dynamics of the meta-parameter. The FIS model also integrates choice and response time to describe internal states and corresponding behavior traits, providing a clear interpretation.

Our results align with prior work showing that learning progress promotes a shift from model-free to model-based control [8, 41], exploitation [21], and choice perseveration [5, 6], consistent across multiple analytical approaches. The competition between model-free and model-based strategies may be modulated by the reliability of their predictions [8, 41]. Learning the task structure facilitates the development of an accurate internal model, leading to reliable predictions from model-based strategies. And the two-step task structure ensures that using the model-based strategy is more advantageous than the model-free strategy [3]. Hence, without overtraining [8], mice did not show a return to model-free control in the later stage of the experiment.

The classical theories suggest that exploitation and exploration form a trade-off, even though exploration is essential for adaptation to change [42, 43, 44]. Strong exploitation generally suppresses exploration, yet frequent block changes in our task necessitate exploration. This study observes that the exploitation is driven by the dominant fast model-based learning. Fast learning and forgetting ensure rapid value updating, which, without affecting the choice stochasticity, promotes the exploitation of the current best option. Such rapid value updating is also observed in the WSLS strategy [2, 45], which can lead to high trial-by-trial variability in choices and, occasionally, suboptimal choices. The mice’s behavior is temporally consistent because the learning and forgetting rates are moderate, the value estimate is not strictly myopic, and the dependency of the choice on value prediction is mediated by a stable internal model. In addition, random exploration, driven by high choice stochasticity, helps avoid being in suboptimal choices.

Choice perseveration reduce cognitive cost and policy complexity [39, 5, 6], but against value-based learning [5, 14] and impede necessary exploration [46, 47]. This study shows that the reduced influence of choice perseveration allows value-based, model-based control to dominate behavior, while the fast choice perseveration provides a compatible fallback when the marginal benefit of model-based planning does not compensate for its cognitive cost [39, 6, 48]. These findings suggest that strategy changes manifest systematically across multiple behavioral axes, with overlaps, supporting the need for integrative approaches beyond narrow analyses of only selected traits.

Adapting to the elevating uncertainty or sudden environmental changes is demanding. Animals and humans usually avoid expending cognitive effort by shifting to a model-free, random strategy [33, 13, 6]. In this study, stay probability and GLMM analysis reveal model-free, exploratory behavior after changes in reward setting or when it is neutral. This pattern might result from two plausible mechanisms. In the neutral block or after a block change, where reward uncertainty is maximal or temporarily unknown, mice may rationally abandon model-based control due to its limited benefit and high cognitive cost, alongside a loss of motivation. However, mice completed a high number of trials per session without showing a lack of motivation. Alternatively, mice continue to rely on value estimates from the preceding block, which mistakenly interact with the current reward settings, yielding behavior that appears exploratory but is actually the result of incomplete adaptation. In a non-neutral block, the reward probability difference provides information to re-establish the knowledge to complete the adaptation. In a neutral block, the lack of information prevents adaptation from completing, so the observed exploratory behavior appears throughout the block. This explanation is also consistent with the observation that mice tend to choose the option that was correct in the preceding blocks.

The RL and FIS modeling might provide insight into differentiating between the two explanations. In these conditions, mice appear to employ a complex model-based strategy with strong fictive updating to optimize their behavior, while being in a suboptimal, choice-repeating state. Yet, the slow forgetting maintains the value estimate learned from the previous block when interpreting the task event, strongly biasing the model-based planning and leading to seemingly exploratory behavior. This interpretation suggests that this exploratory behavior may be driven by the interaction between previously learned value estimates and current events, an incomplete adaptation that later can be resolved in a non-neutral block but not in a neutral one. Under high task demands, both avoidance of and reliance on complex strategies are possible [33, 34]. This study mainly observes reliance on complex strategies, possibly because the task demands do not exceed the mice’s cognitive limits. An implication for further examination is that internal self-persistence may fundamentally drive attempts at model-based planning based on incomplete adaptation, rather than passively abandoning the task.

Besides, the FIS model reveals the discrete strategy switch in the trial resolution. A reward reinforces the individual action and also the applied strategy, especially the optimal one. It implies a generalized role of reward in modulating meta-learning. How obtaining the reward reinforces the single actions is well characterized in the RL framework [49]. However, the reward signals [49, 50] and value expectation [51, 52, 53] are represented in distributed neural circuits and might modulate neural activity across a wide range of brain regions. Hence, reward signals might play a more general role in mediating value-based behavior, consistent with the findings here. Then, one potential function of the brain-wide reward signals might be to support the implementation of behavioral strategy regulation.

This research suggests that discrete switches and continuous adaptation may be parallel processes operating on different timescales, with potential interactions. Previous frameworks often assume continuous or discrete strategy changes. Which view to take often depends on the influencing factors being investigated. Research focusing on the long-term effects of learning progress and environmental features usually assumes a continuous strategy adaptation [7, 19, 2, 15]. Studies highlighting the influence of internal state often hypothesize the discrete strategy switch [16, 18]. In complement to previous studies, our research shows evidence of how two processes co-occur and interact. Besides, Ashwood et al. [16] shows that the discrete HMM-based model captures the mice’s perceptual decision-making behavior better than the continuous model. This finding might be partially due to the fact that the behavior from the learning phase is not included in the model comparison, whereas this study suggests that the most prominent strategy adaptation occurs alongside learning on the long-time scale.

At the methodological level, the FIS model associates different behavioral patterns with distinct response patterns, which together suggest cognitive strategies and the underlying internal state. Behavior observed in the experiment is often compressed into individual binary responses, losing details necessary for interpreting the underlying cognitive process, whereas the response (reaction) time provides a fine-grained measure of the ongoing cognitive activities. Modeling reaction time often uses the sequential sampling models, which traditionally assume that response times across trials follow the same distribution [54, 55, 56]. On the one hand, previous attempts at jointly modeling reaction time and value-based choice have mainly focused on integrating RL with the evidence accumulation model [57, 58, 59, 60, 61]. These models provide reaction-time modeling with learning dynamics by leaving temporal variability to the RL mechanism. The FIS model demonstrates another direction by simultaneously considering the different patterns of choice and reaction time, along with their correspondence, without imposing assumptions about the learning process.

This study also has its limitations. A systematic examination of strategy switching requires a complex behavior task and a complex cognitive model. Although our dynamic modeling approaches extract the dynamics, we refrain from interpreting the raw trajectory directly, as it may contain noise and fluctuations due to inference difficulties, despite a few exploratory analyses suggesting there is a latent temporal structure in meta-parameter dynamics. While acknowledging this limitation, we call for further investigation of the meta-parameter dynamics extracted from mice behavior performing simpler tasks with a simpler model.

Among several other promising directions for future work, we particularly advocate for investigating the neural substrates underlying discrete switches and continuous adaptation in behavioral strategy. The continuous adaptation relies on its past trajectories and current sensory inputs, much like a recurrent neural network, which might take place in the prefrontal cortex [25]. A discrete switch, if within a finite set of hidden states, requires explicit memory of those states. Hence, it might be implemented via interactions between the hippocampus and the basal ganglia. A lesion study of the hippocampus or the prefrontal cortex might help separate the neural substrates of the two processes. Moreover, developing algorithmic frameworks that formally integrate discrete and continuous components could yield candidate models for a flexible intelligent system with meta-learning capability.

## 4 Methods

### 4.1 Animals

All experimental procedures were approved by and performed in accordance with guidelines from the Okinawa Institute of Science and Technology Experimental Animal Committee. Ten Tph2-tTA male mice were used for the behavioral experiment. All mice were housed separately at 24^*o*^*C* with a 12 h light-dark cycle (lights on 7 am -7 pm). The training began at 8-14 weeks of age and was conducted during the light cycle, 6 days a week. Mice were water-deprived 24 h before the beginning of the training and received water only in the experiment as a reward (∼ 800 ul). Water deprivation was temporarily removed for 60 minutes once the animals’ body weight dropped by more than 15% during the experiment. The animal received 1 hour of ad lib water access on the day off.

### 4.2 Behavioral apparatus

The experiment was conducted in the 12×12 cm operant boxes controlled by pyControl [62]. Five ports with LED lights were located on the front wall of the boxes for nose-poking, stimuli delivery, and water administration. The center port was for trial initiation. The choice ports were located 4 cm to the left and right of the center port, and the two-state ports were positioned 1.6 cm above and below the center port. Two state ports were connected with two solenoids for water delivery. Solenoids were calibrated monthly to ensure the accuracy of the water delivered. A speaker above five ports delivered the auditory cues.

### 4.3 Behavioral task and training protocol

When the trial was ready to start, the center port was illuminated. The trial began after nose-poking at the center port, where choice ports were illuminated. In the forced-choice trials, in which animals were forced to choose a specific option, only the randomly specified port lit up. Two-choice ports were illuminated during the free trial, with two options available. 75% of the trials are free trials in the task. The first-step choice led to either of the two states being determined by the transition matrix, which was counterbalanced between animals but fixed within the subject. In transition type A, the left choice commonly (80%) led to the up state but rarely (20%) to the down state. Type B transition had a reversed structure.

After a 200 ms delay following the first step choice, the state port lit up with a 1000 ms auditory stimulus. The stimulus was either a 5 or 12 kHz tone for up or down states, or vice versa (counterbalanced between animals). After presenting the tone, animals were allowed to nose-poke the active state port to confirm the state. After a 200 ms delay following state check, a 500 ms outcome cue was delivered (10 kHz tone signaling reward delivery and the white noise signaling the reward omission). After the outcome delivery, there was a 2-4 s inter-trial interval (ITI).

The reward probabilities of the two states changed in an anti-correlated manner between blocks. There were three block types: up block (reward probability of up state is 80%, 20% for down state), down block (20% for up state and 80% for down state), and neutral block (50% for both). In non-neutral blocks, block changes happened after 5-15 trials once the exponential moving average of the correct response (i.e., choosing the option that commonly leads to a state with higher reward probability) in the last eight free trials reached 75%. In the neutral block, the block changed after 20-30 trials.

The training protocol was the same as described in [36]. The training was divided into several phases. In phases 1-3, animals were instructed to build up the action sequence of nose-poking the center, choice, and state ports and learn the transition matrix. In phase 4 and sub-phases, animals were instructed to learn the reward probability and its changing rule. In phase 1.1, the state ports were randomly lit, and nose-poking the active port always led to a reward. Once animals completed 50 or more trials per session, training proceeded to phase 1.2, where state cues were introduced. After performing more than 50 trials per session, phase 2 started. In phase 2, either the left or right choice port was active, and following the transition, the state port was activated. Once the same criteria were met, animals were trained in phase 3, where the initiation step was introduced. After phase 3, animals could perform the whole action sequence and learn the transition matrix. Once the animal could perform more than 75 trials per session, phase 4 began. In sub-phases 4.1 - 4.5, we gradually increased the proportion of free choice and introduced the reward probability and performance-dependent block-changing rule. Starting from phase 4.6, 75% of trials were free-choice, and the block changed based on performance. The training schedule changed from two 45-minute sessions to one 90-minute session per day. In phase 4.6, the reward amount gradually decreased from 12 ul to 4 ul to encourage the animal to perform more trials. The task is the same as described before, except that the reward probabilities were 10%, 90%, and 50%. The animal that performed 5-6 block one session was transitioned to the final phase 4.7, with the full task. In this phase, performance was considered stable once the animal completed more than 6 blocks on 3 consecutive days. The analysis included data from phase 4.5 to 4.7, which are named stage I to III, respectively.

**Table 1:** Experiment condition. In stage I, the reward size decreased gradually, while the reward probability settings (e.g., 10% and 90%) were highly discriminative to motivate and facilitate learning. In stages II and III, the reward size was fixed as 4ul, and the reward probability settings became less distinguishable (e.g., 20% and 80%). The animals were considered to be in stage III only when they showed stable performance across more than 6 blocks on two consecutive days. Since the experiment length is fixed and the block change depends on the performance in a non-neutral block, performing more blocks indicates a higher choice accuracy.

| Stage | Reward size (ul) | Reward settings (bad/good/neutral) | Complete criterion | # of blocks | # of trials | Accuracy ** |
| --- | --- | --- | --- | --- | --- | --- |
| I | 10 ~ 4 | 10%, 90%, 50% | $\geq 5$ blocks (for 1 days) | 289 | 14961 | 48.05% |
| II | 4 | 20%, 80%, 50% | $\geq 6$ blocks (for 2 days) | 325 | 14779 | 56.83% |
| III | 4 | 20%, 80%, 50% | NA | 2181 | 74888 | 64.95% |

\*\* Percentage of correct response in the non-neutral block. Because the reward settings change with performance, the accuracy is upper-bounded.

### 4.4 Statistical analysis

All behavioral analyses were performed using custom Python, R, and MATLAB scripts with publicly available packages. The packages used will be specified in the following sections. The statistical tests were selected based on the data distribution. For normally distributed data, we use paired *t*-tests when conducting within-subject comparisons with equal sample sizes and unpaired *t*-tests otherwise. For datasets that do not follow a normal distribution, we used the Wilcoxon signed-rank test and the Mann-Whitney U-test, respectively.

#### 4.4.1 Task events, behavioral measure, and response times

All task events, including the choices, transition types, and outcomes, were extracted from the experiment records. Due to the nature of the task, instead of analyzing the actual choice (i.e., left and right), the analysis focuses on the stay/switch behavior (i.e., whether the current choice is the same as the last one).

Four response measures were derived from the response time, including early response, initiation time, decision time, and latency to cue. Early Response is defined as a binary variable indicating whether the animal nose-poked at the center poke to initiate the trial during the ITI. If the animal made an early response, the trial would initialize only if the animal withdrew from the port and redid the nose-poking after ITI, which tended to take longer. To assess the initiation time without bias, it is defined as the time to nose-poke the center poke since the end of ITI for valid responses or since the end of the trial for early responses. Similarly, decision time is defined as the time to make the first-step choice, regardless of whether the choice is valid (invalid choices are defined as nose-poking the inactive poke in forced-choice trials). During the delay between the first-step choice and second-step state presentation, the animal always tended to nose-poke the illuminated state port. Besides, animals rarely chose the inactive port, ignoring the cue. Latency to cue is defined as the time to confirm the second-step state for valid and invalid responses, but not the cue-independent response. The latter three measures were in seconds, and the between-subject comparison was based on their median value, unless it is specified otherwise.

### 4.4.2 Generalized linear mixed model (GLMM)

Multiple variables may affect the behavior and response time measures, which exhibit high individual variability in the task. To examine those effects without interference, we built several GLMMs using the fitglme function in MATLAB 2023b. The link function was chosen based on the distribution of the dependent variable, while all other configurations retained their default settings. All included variables were assumed to vary between subjects, and a random effects matrix with subjects as grouping factors was included. For the between-group comparison of the main effects, we construct a separate model with the specified model structure and the interaction term between the grouping factors (e.g., learning stage) and regressors in the fixed and random effect matrices. The grouping factors were also included as regressors in fixed and random effect matrices. The interaction term shows whether there is a significant difference in the main effect between groups. The main effect of grouping factors indicates whether the difference in baseline (i.e., the intercept) is significant between groups. In GLMM, including the interaction term, the main effect is conditional on the grouping factor and does not reflect the overall trend. Hence, we interpret the main effect in the version without interaction terms in the main text.

Seven GLMMs were constructed to predict the stay/switch behavior in free trials using the logit link function. The several model formulas were:

1. stay/switch ∼ intercept + choice + outcome + transition + outcome × transition + (variables | subject)
2. stay/switch ∼ intercept + choice + outcome + transition + correct + outcome × transition + (variables | subject)
3. stay/switch ∼ intercept + trials + choice + outcome + transition + correct + outcome × transition + (variables | subject)
4. stay/switch ∼ intercept + trials + choice + ∆value + transition + outcome × transition + (variables | subject)
5. stay/switch ∼ intercept + trials + choice + ∆value + entropy + transition + outcome × transition + (variables | subject)
6. stay/switch ∼ intercept + choice + ∆value + entropy + transition + outcome × transition + (variables | subject)
7. stay/switch ∼ intercept + choice + ∆value + transition + outcome × transition + (variables | subject) The stay/switch was coded as 1 if the animal made the same choice as in the last trial and 0 otherwise.

The regressors were coded as:

- trial: the trial index starting from 1.
- choice: previous action *a*_*t*−1_, 0.5 if the previous choice was left, -0.5 otherwise.
- outcome: previous outcome *r*_*t*−1_, 0.5 if the previous trial was rewarded, -0.5 if omitted.
- transition: 0.5 if the previous transition is common, -0.5 if rare.
- correct: in the non-neutral block, 0.5 if the previous choice was correct, -0.5 otherwise. In the neutral block, due to the task setting, this variable is always 0.5.
- outcome × transition: 0.5 if the previous trial was reward-common transition or omission-rare transition, -0.5 otherwise.
- ∆value: inferred value difference. The animal’s estimate of the difference in reward probabilities between the chosen and unchosen options prior to the current trial.

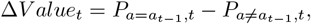

where,

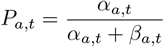

where,

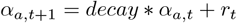

and,

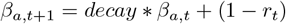

where *decay* = 0.5.

- Entropy: the entropy of the chosen option in the last trial.

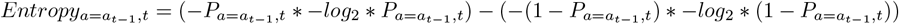

The GLMM model for response time analysis was adopted from the fourth version with some modifications.

- Early Response ∼ intercept + time + ∑ outcome + neutral + late phase + ∆Value(-1) + transition(-1) + ITI + stay (free) + (variable | subject)
- initiation Time ∼ intercept + time + ∑ outcome + neutral + late phase + ∆Value(-1) + transition(-1) + ITI + early + stay (free) + (variable | subject)
- Decision time ∼ intercept + time + ∑ outcome + neutral + late phase + ∆Value(-1) + transition(-1) + ITI + invalid + free trial + stay (free) + stay (force) + (variable | subject)
- Latency to Cue ∼ intercept + time + ∑ outcome + neutral + late phase + ∆Value(-1) + transition(-1) + ITI + cue-independent + free trial + transition + state (same) + stay (free) + stay (force) + (variable | subject)

The included regressors fall into five categories.

1. Long-term effects:
  - time: time elapsed since the start of the experiment in seconds.
  - ∑ outcome: the number of rewards obtained so far.
2. Block-wise effects:
  - neutral: 1 for trials in the neutral block, 0 otherwise.
  - late phase: 0 for the first 10 trials after the block changes, 1 otherwise.
3. Last-trial effects:
  - ∆value(-1): inferred value difference in the last trial.
  - transition (-1): the transition type in the last trial.
  - ITI: Inter-trial interval before the current trial, 3-5 seconds.
4. Current-trial effects:
  - early: 1 if the early response has been made, 0 otherwise.
  - invalid: 1 if the invalid choice has been made in forced-choice trials, 0 otherwise.
  - cue-independent: 1 a cue-independent response has been made, 0 otherwise.
  - free: 1 for free trials, 0 otherwise.
  - transition: the transition type in the current trial.
5. Effects of the contrast between the last and current trial:
  - state (same): 1 if the states presented in the last and current trial are the same, 0 otherwise.
  - stay (free): 1 if the current choice is the same as in the last trial in the free-choice trial, 0 otherwise.
  - stay (stay): 1 if the current choice is the same as in the last trial in the forced-choice trial, 0 otherwise.

Some regressors show differences only in part of the trials; for example, invalid choice is presented only in forced-choice trials. To estimate the effect of those regressors without bias, we construct a separate model using a subset of the dataset with the same structure. Specifically, we only include the forced-choice trials to estimate the invalid and stay (force), and only free-choice trials to estimate the stay (free).

#### 4.4.3 Finite internal state model (FIS)

The presented finite internal state model extends the hidden Markov model in two aspects. The FIS model assumes agents are in transit between multiple hidden states. The transition probabilities between states are determined by the last action taken, the outcome observed, and the transition type experienced (i.e., common or rare) in the last trials, similar to the previous study [22]. And there are multiple observations in each state. Each internal state possesses a distinct emission probability for stay/switch behavior and response-time measures, indicating the strategy used and the corresponding internal state. To capture the overall trend across subjects *i* and sections *j*, we constructed the model hierarchically, assuming a subject-level random intercept that affects the transition and emission probabilities.

FIS models has *N* hidden state *s, s* ∈ 1, 2, …, *N*. The probabilities of being at those states at the initial time *t* = 0, the initial distribution, are,

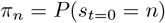

with the constraint that, 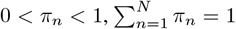.

The subject (i.e., mouse) *i* transits between states according to the transition probability. The transition probability from state *m* to *n* is parameterized as the linear combination of fixed and random effects *η*_*n,m*_ with the softmax function, taking state 1 as the baseline,

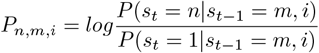

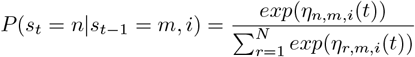

where,

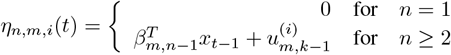

where, 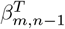 is the fixed regression coefficients for transition probability. *x*_*t*−1_ are the inputs (i.e., outcome, transition type, and choice of the last trial) that affect the transition probability 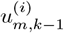 is the subject-level random effect, which is implemented by non-centered parameterization,

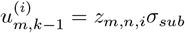

where,

*z*_*m,n,i*_ is sampled from 𝒩 (0, 1) and scaled by the subject-level deviation *σ*_*subj*_.

Each state has its own emission probability of five observations, including two binary variables (i.e., stay/switch choice and early response) and three continuous variables (i.e., response vigor, decision time, and latency to cue),

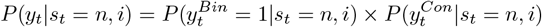

where,

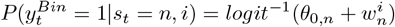

where, *θ*_0,*n*_ is the intercept for emission probability (i.e., baseline), 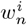 is the subject-level random effect that computed by the same non-centered parameterization.

Since three continuous variables are derived from response times, we model them by an exponential Gaussian distribution [54],

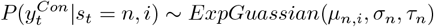

where,

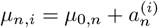

where, *µ*_*n,i*_ is the mean of the distribution, which is the sum of the baseline *µ*_0,*n*_ and the subject 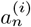 random mean. *σ*_*n*_ and *τ*_*n*_ are fixed for simplicity.

Instead of using the Expectation-maximization algorithm, we implement a forward algorithm with the MCMC sampling method, which shares the identical core concept but is easier to implement in Rstan.

#### 4.4.4 RL Strategy Models

We construct different RL learning strategy models for simulation and model fitting. Base models can be categorized into three types: model-free, model-based, and hybrid models. There are five additional rules to describe other phenomena found in the relevant literature, including loss aversion, asymmetric learning [36], forgetting, choice perseveration, and fictive learning [40].

In the task, the agent chooses an action *a* ∈ (*left, right*), which leads to the second-step state *s* ∈ (*up, down*), where the outcome *r* ∈ (0, 1) is delivered. Agents update the action value *Q*(*a*) based on past experience through various learning strategies, yet the action selection always follows the softmax function.

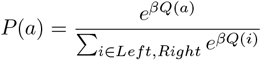

The model-free category encompasses the model-free model with eligibility trace, MF-lambda, as well as its two specific variants: direct model-free, MF, and model-free with memory strategy, MF-memory. The MF-lambda agent updates its action value of chosen options *Q*_*mf*_ (*a*) and state value of experienced state *V* (*s*) as follow,

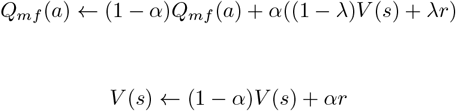

The MF-lambda has two meta-parameters, learning rate (*α*) and eligibility trace (*λ*). The model is termed MF and MF-memory when *λ* is 1 or 0, respectively.

The model-based category includes the model-based (MB) and latent-state inference (LSI). In model-based strategies, agents exploit the learned world model, the transition matrix between actions and state (*P* (*s*|*a*)). The MB agent learns the state value as follows,

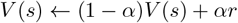

and then compute the action value by,

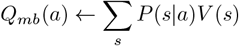

Similarly, the agent might assume that there are two latent states in which either of the two second-stage states is better and update the beliefs of being one latent state (*h* ∈ *h*_*up*_, *h*_*down*_) using Bayesian inference [36].

Specifically, the agent estimates the *P* (*h*_*up*_) by,

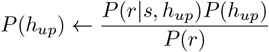

where, the likelihood *P* (*r*|*s, h*_*up*_) is assumed known to agents 2.

**Table 2:** Likelihood table.

| (a) $P(r\vert s, h_{up})$ | | | (b) $P(r\vert s, h_{down})$ | | |
| --- | --- | --- | --- | --- | --- |
|  | Reward | Omission |  | Reward | Omission |
| Up | 0.8 | 0.2 | Up | 0.2 | 0.8 |
| Down | 0.2 | 0.8 | Down | 0.8 | 0.2 |

The marginal likelihood *P* (*r*) is calculated as,

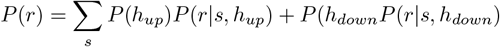

Since the reward settings vary in the task, the agent might assume that the environment would change with a certain probability *ψ*. Hence, the posterior is updated as,

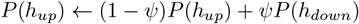

Therefore, the state values are updated as,

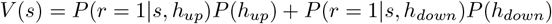

and the action value is updated as in MB.

A hybrid model consists of multiple model-free and model-based strategies, and the action value is computed as,

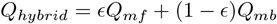

It provides six hybrid models combining three model-free strategies with two model-based strategies (MF-MB, MF-lambda-MB, MF-memory-MB, MF-LSI, MF-lambda-LSI, MF-memory-LSI).

We also incorporate additional rules based on the above models. Firstly, the agent may develop an aversion to reward omission to implement the WSLS strategy (Loss). Specifically, reward omission is −*κ* rather than 0, which captures the strength of loss aversion.

A similar strategy is for the agent to learn asymmetrically from rewards and omissions (Asym). In models other than the LSI, it is implemented with two learning rates: *α*_*pos*_ for rewards and *α*_*neg*_ for omissions. The LSI model does not explicitly have the learning rate parameter, so the asymmetric rule is implemented by modifying the likelihood table as in 3,

**Table 3:** Likelihood table (asymmetric)

| (a) $P(r\vert s, h_{up})$ | | | (b) $P(r\vert s, h_{down})$ | | |
| --- | --- | --- | --- | --- | --- |
|  | Reward | Omission |  | Reward | Omission |
| Up | 0.4 | 0.5 | Up | 0.1 | 0.5 |
| Down | 0.1 |  | Down | 0.4 |  |

So, the agent takes the omission in both states as the same observation.

The action and state value might be forgotten if it is not chosen *Q*(*a*_−_) and unvisited *V* (*s*_−_) (Forget). The forgetting rule is implemented as,

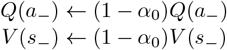

Choice perseveration encourages choice repetition over time. The choice perseveration rule (Perse) assumes that agents maintain a choice kernel (CK) that reflects the choice history,

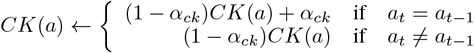

The choice perseveration contributes to the action value by,

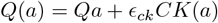

The last additional rule is fictive learning (FL), which updates the value of an inexperienced event (i.e., an unvisited state and an unchosen option) by exploiting the learned task structure between the inexperienced and experienced events. In MF-lambda model, the updating rule for state and action value can be rewritten as,

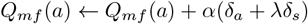

and

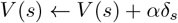

where *δ*_*a*_ = *V* (*s*) − *Q*_*mf*_ (*a*) and *δ*_*s*_ = *r* − *V* (*s*) are the prediction error for action and state value.

The agent might use the prediction error from experienced events to update the unchosen action and unvisited state by their inferred correlation between them *η*_*a*_ and *η*_*a*_ by,

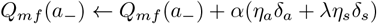

and

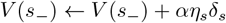

Both correlation factors are zero when the agent believes the two actions and states are changing independently, negative if they are anticorrelated, and positive if they are changing in the same direction. In model simulation and fitting, we simplified the model in two aspects. Since the transition matrix was fixed and well-instructed, we assume that the agent believes the correlation between state and action is the same (i.e., *η*_*s*_ = *η*_*a*_ = *η*). For the same reason, we assume that the agent already learns the correlation during the training and *η* is a hyperparameter that does not depend on other behavior measures (e.g., the magnitude of *δ*_*a*_ and *δ*_*s*_).

#### 4.4.5 Model Simulation

To examine the analysis tool for strategy discovery, we simulate several learning strategies that are commonly presented in the relevant task. The agent’s included parameters and settings are described in 4. To make sure the model simulation result is not limited to specific parameter settings, we add a noise term *noise* ∼ 𝒩 (0, 0.05 × |*parameter*|) for each agent. For each model, we simulated 14 agents over 20 sessions, with each session comprising 400 trials, a typical length in animal experiments.

**Table 4:** Model and parameter settings.

| Model | $\beta$ | $\alpha$ | $\alpha_0$ | $\lambda$ | $\kappa$ | $\alpha_{cp}$ | $\epsilon_{cp}$ | $\psi$ | $\eta$ | $\epsilon$ |
| --- | --- | --- | --- | --- | --- | --- | --- | --- | --- | --- |
| MF | 3 | 0.4 | 0 | 1 |  |  |  |  |  |  |
| MF-memory | 3 | 0.4 | 0 | 0 |  |  |  |  |  |  |
| MF-Loss | 3 | 0.4 | 0 | 1 | 0.7 |  |  |  |  |  |
| MF-Prese | 3 | 0.4 | 0 | 1 | 0 | 0.4 | 0.5 |  |  |  |
| MB | 3 | 0.4 | 0 |  |  |  |  |  |  |  |
| LSI | 3 |  |  |  |  |  |  | 0.8 |  |  |
| MF-MB | 3 | 0.4 | 0 | 1 |  |  |  |  |  | 0.5 |
| MF+LSI | 3 | 1 | 0 | 1 |  |  |  | 0.8 |  | 0.5 |
| MB-FL | 3 | 0.4 | 0 |  |  |  |  |  | 1 |  |
| Random | 0.01 | 0.4 | 0 | 1 |  |  |  |  |  |  |

Abbreviation of model names. MF: direct model-free; MF-memory: model-free with memory; MF-Loss: model-free with loss aversion; MF-Prese: model-free with choice perseverance; MB: model-based; LSI: latent-state inference; MF-MB: the hybrid model of direct model-free and model-based; MF+LSI: the hybrid model of direct model-free and latent-state inference; MB-FI: model-based with fictive learning; random: random selection.

#### 4.4.6 Bayesian Model fitting

We implement the Bayesian fitting with Rstan 2.32.6 with 4 MCMC chains for 2000 iterations (500 warm-up runs). We first conducted extensive model fitting to identify potential learning strategies. We included base models: three MF and two MB models, as well as six hybrid models, each with five additional learning rules. A few models were not implemented due to conflicts in learning rules, resulting in 61 models for the first step. The model fitting was performed for each subject and session to account for high subject- and session-level variability. Note that the model fit in this and the next session included both free- and forced-choice trials for variable updating. But the total likelihood is the sum of the choice probabilities in free-choice trials only. The Bayesian Information criterion (BIC) score is used for model comparison. Specifically, we compute the Δ BIC score as the difference between the BIC score of the best-fitting model and the BIC score of the model being evaluated.

#### 4.4.7 Forward particle filtering

We improve the particle filtering [19, 2] to filter the time-varying meta-parameters in RL from the choice by including an adaptive forward step in evaluating the important weight of particles inspired by the auxiliary particle filter [63]. This particle filtering is a sequential Bayesian inference approximation using the Monte Carlo method, comprising three steps: prediction, resampling, and propagation. In standard particle filtering [19, 2] for a given RL model, ℛℒ, for particle *i* at trial *t*, define,

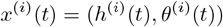

where *h*^(*i*)^(*t*) and *θ*^(*i*)^(*t*) represent the hidden variables (e.g., action value, *Q*_*mf*_, *Q*_*mb*_; state value, *V*) and meta-parameters, (e.g, inverse temperature, *β*; learning rate, *α*), respectively, to predict the choice, *a*(*t*) according to ℛℒ. The particles are then weighted by their predicted probability of the actual choice for resampling: particles with high weights are preserved and replicated. Then the resampled particles are propagated to the next by the RL model for *h*^(*i*)^(*t*) and by a random walk for *θ*^(*i*)^(*t*). The aim is to estimate the dynamics of meta-parameters (i.e., the posterior distribution of *θ*(*t*)), by grid searching the hyperparameter, *ψ*, (i.e., the noise size in random walk, *σ*, and initial distribution, *P* (0), and the RL models) that maximizes the predicted probability of the actual choice.

At initialization, assuming that hidden variables and meta-parameter are independent, N independent initial particles *x*^(*i*)^(0) are sampled from the prior distribution *P* (0) = *p*(*x*_0_)*p*(*θ*_0_)by,

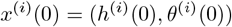

where,

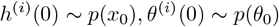

And the particle’s importance weights, *w*^(*i*)^(*t*), are initialized so that,

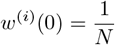

Thereafter, in each trial *t*, each particle predicts the probability of choosing the left option, *P*_*L*_(*t*), computed by elements in the particles with the RL model. *R*_*L*_(*t*) is transformed as the particle’s importance weight, *w*(*t*). For each particle *i*,

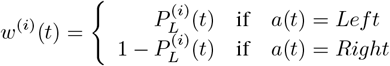

*w*^(*i*)^(*t*) is then normalized to 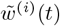 so that 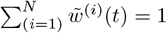. When the effective sample size (ESS), *ESS*(*t*),

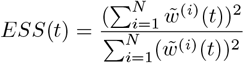

is below an adaptive threshold, *ξ*, resampling take place based on the 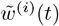. Here, the algorithm is illustrated with systematic resampling [64]. Particles are aligned on an interval [0, 1] and divided into sections with length equal to 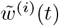. Then the logarithm randomly picks the first particles from the interval 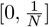, followed by the other particles picked with equidistant intervals of ^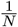^ between two particles. This method ensures that particles with higher 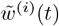 are preserved and replicated. The posterior distribution of meta-parameters, given the observation (action, *a*(1 : *t*); state, *s*(1 : *t*); and reward, *r*(1 : *t*)) so far, is approximated as,

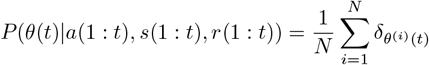

where the *δ*_(.)_ is the Dirac function. Those particles will then be propagated. For meta-parameter *j* in *θ*^(*i*)^(*t*), it is updated by random walk,

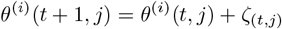

where *ζ*_(*t,j*)_ is drawn independently drawn from 𝒩 (0, *σ*_(*j*)_). And *h*^(*i*)^(*t*) is updated based on the ℛℒ and *θ*(*t* + 1),

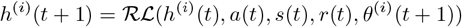

which then served as the prior distribution for the filtering in the next trial, *t* + 1.

The aim is to estimate the dynamics of meta-parameters (i.e., the posterior distribution of *θ*(*t*)), by finding the hyperparameters, *ψ*, (i.e., the noise size in random walk, *σ*, and initial distribution, *P* (0)), given the particular ℛℒ, that maximizes the predicted probability of the actual choice.

The original weighting and resampling algorithm relies solely on the current observation, potentially leading to particles with extreme predictions being given higher weights, regardless of the actual, unknown choice probability. To overcome this, we evaluate choice probability using both the current and the adjacent future trials, termed the forward step. The adaptive forward step uses future observations to preselect particles likely to generate them. After calculating the *w*(*t*), we update the *h*(*t*) to predict the *a*(*t* + 1),

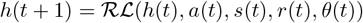

where the meta-parameters are those at trial *t* without random walking. The same weighting step yields the 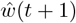, the forward weight at trial *t* + 1. Then the forward step continues to trial *t* + 2. The forward step stops when it reaches the predetermined, maximal steps or 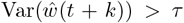, representing the idea that the current trial provides enough information to differentiate the predictability of each particle. Therefore, a weight matrix 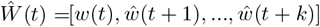 is obtained. Since the choice probability is more likely to change in the distal future, the importance of the forward weight in the far future is discounted.

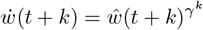

where,

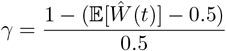

reflecting the idea that the temporal discounting factor depends on the predictability of all particles. The distal observation matters more when the particles are generally far from the true region. The overall high accuracy leads to steep temporal discounting, so that evidence from recent trials is emphasized for their temporal relevance when the current meta-parameter distribution is likely to be in the true region.The 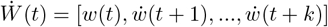 is then normalized for the resampling of the original particle *x*(*t*) = [*h*(*t*), *θ*(*t*)].

In the current study, we utilize the Optuna framework [65] to effectively search for the hyperparameters. We conduct 1,000 search trials with 5,000 particles each to cover the possible ranges of hyperparameters. *τ* is set as 0.05.

## References

[1] John P O’Doherty, Jeffrey Cockburn, and Wolfgang M Pauli. Learning, reward, and decision making. Annual review of psychology, 68:73–100, 2017.

[2] Makoto Ito and Kenji Doya. Validation of decision-making models and analysis of decision variables in the rat basal ganglia. Journal of Neuroscience, 29(31):9861–9874, 2009. ISSN 0270-6474. doi:10.1523/JNEUROSCI.6157-08.2009. URL https://www.jneurosci.org/content/29/31/9861.

[3] Thomas Akam, Rui Costa, and Peter Dayan. Simple plans or sophisticated habits? state, transition and learning interactions in the two-step task. PLoS computational biology, 11(12):e1004648, 2015.

[4] Robert C Wilson and Anne GE Collins. Ten simple rules for the computational modeling of behavioral data. eLife, 8:e49547, nov 2019. ISSN 2050-084X. doi:10.7554/eLife.49547. URL 10.7554/eLife.49547.

[5] Kevin J Miller, Amitai Shenhav, and Elliot A Ludvig. Habits without values. Psychological review, 126(2):292, 2019.

[6] Samuel J Gershman. Origin of perseveration in the trade-off between reward and complexity. Cognition, 204: 104394, 2020.

[7] Nicholas A. Roy, Ji Hyun Bak, Athena Akrami, Carlos D. Brody, and Jonathan W. Pillow. Extracting the dynamics of behavior in sensory decision-making experiments. Neuron, 109(4):597–610.e6, 2025/12/02 2021. doi:10.1016/j.neuron.2020.12.004. URL 10.1016/j.neuron.2020.12.004.

[8] Nathaniel D Daw, Yael Niv, and Peter Dayan. Uncertainty-based competition between prefrontal and dorsolateral striatal systems for behavioral control. Nature Neuroscience, 8(12):1704–1711, 2005. ISSN 1546-1726. doi:10.1038/nn1560. URL 10.1038/nn1560.

[9] Kevin S. Chen, Anuj K. Sharma, Jonathan W. Pillow, and Andrew M. Leifer. Navigation strategies in caenorhabditis elegans are differentially altered by learning. PLOS Biology, 23(3):1–24, 03 2025. doi:10.1371/journal.pbio.3003005. URL 10.1371/journal.pbio.3003005.

[10] Lenca I Cuturela and Jonathan W Pillow. Internal states emerge early during learning of a perceptual decision-making task. bioRxiv, Sep 2025. ISSN 2692-8205 (Electronic); 2692–8205 (Linking). doi:10.1101/2024.11.30.626182.

[11] Cooper D. Grossman, Bilal A. Bari, and Jeremiah Y. Cohen. Serotonin neurons modulate learning rate through uncertainty. Current Biology, 32(3):586–599.e7, 2025/12/02 2022. doi:10.1016/j.cub.2021.12.006. URL 10.1016/j.cub.2021.12.006.

[12] Wouter Kool, Fiery A. Cushman, and Samuel J. Gershman. When does model-based control pay off? PLOS Computational Biology, 12(8):1–34, 08 2016. doi:10.1371/journal.pcbi.1005090. URL 10.1371/journal.pcbi.1005090.

[13] Wouter Kool, Samuel J. Gershman, and Fiery A. Cushman. Planning complexity registers as a cost in metacontrol. J. Cognitive Neuroscience, 30(10):1391–1404, October 2018. ISSN 0898-929X. doi:10.1162/jocn_a_01263. URL 10.1162/jocn_a_01263.

[14] Lucy Lai and Samuel J. Gershman. Human decision making balances reward maximization and policy compression. PLOS Computational Biology, 20(4):1–32, 04 2024. doi:10.1371/journal.pcbi.1012057. URL 10.1371/journal.pcbi.1012057.

[15] Yoav Ger, Eliya Nachmani, Lior Wolf, and Nitzan Shahar. Harnessing the flexibility of neural networks to predict dynamic theoretical parameters underlying human choice behavior. PLOS Computational Biology, 20(1):1–22, 01 2024. doi:10.1371/journal.pcbi.1011678. URL 10.1371/journal.pcbi.1011678.

[16] Zoe C. Ashwood, Nicholas A. Roy, Iris R. Stone, Anne E. Urai, Anne K. Churchland, Alexandre Pouget, Jonathan W. Pillow, and The International Brain Laboratory. Mice alternate between discrete strategies during perceptual decision-making. Nature Neuroscience, 25(2):201–212, 2022. doi:10.1038/s41593-021-01007-z. URL 10.1038/s41593-021-01007-z.

[17] Scott S. Bolkan, Iris R. Stone, Lucas Pinto, Zoe C. Ashwood, Jorge M. Iravedra Garcia, Alison L. Herman, Priyanka Singh, Akhil Bandi, Julia Cox, Christopher A. Zimmerman, Jounhong Ryan Cho, Ben Engelhard, Jonathan W. Pillow, and Ilana B. Witten. Opponent control of behavior by dorsomedial striatal pathways depends on task demands and internal state. Nature Neuroscience, 25(3):345–357, 2022. doi:10.1038/s41593-022-01021-9. URL 10.1038/s41593-022-01021-9.

[18] Adam J. Calhoun, Jonathan W. Pillow, and Mala Murthy. Unsupervised identification of the internal states that shape natural behavior. Nature Neuroscience, 22(12):2040–2049, 2019. doi:10.1038/s41593-019-0533-x. URL 10.1038/s41593-019-0533-x.

[19] Kazuyuki Samejima, Kenji Doya, Yasumasa Ueda, and Minoru Kimura. Estimating internal variables and paramters of a learning agent by a particle filter. In S. Thrun, L. Saul, and B. Schölkopf, editors, Advances in Neural Information Processing Systems, volume 16. MIT Press, 2003. URL https://proceedings.neurips.cc/paper_files/paper/2003/file/db60b95decdeed944b4cd8685417cfdc-Paper.pdf.

[20] Nhat Minh Le, Murat Yildirim, Yizhi Wang, Hiroki Sugihara, Mehrdad Jazayeri, and Mriganka Sur. Mixtures of strategies underlie rodent behavior during reversal learning. PLOS Computational Biology, 19(9):1–28, 09 2023. doi:10.1371/journal.pcbi.1011430. URL 10.1371/journal.pcbi.1011430.

[21] Sarah Jo C Venditto, Kevin J Miller, Carlos D Brody, and Nathaniel D Daw. Dynamic reinforcement learning reveals time-dependent shifts in strategy during reward learning. elife, December 2024. doi:10.7554/elife.97612.2. URL 10.7554/eLife.97612.2.

[22] Makoto Ito and Kenji Doya. Parallel representation of value-based and finite state-based strategies in the ventral and dorsal striatum. PLoS computational biology, 11(11):e1004540, 2015.

[23] Sebastian Thrun and Lorien Pratt. Learning to learn: Introduction and overview. In Learning to learn, pages 3–17. Springer, 1998.

[24] Kenji Doya. Metalearning and neuromodulation. Neural Networks, 15(4):495–506, 2002. ISSN 0893-6080. doi:10.1016/S0893-6080(02)00044-8. URL https://www.sciencedirect.com/science/article/pii/S0893608002000448.

[25] Jane X. Wang, Zeb Kurth-Nelson, Dharshan Kumaran, Dhruva Tirumala, Hubert Soyer, Joel Z. Leibo, Demis Hassabis, and Matthew Botvinick. Prefrontal cortex as a meta-reinforcement learning system. Nature Neuroscience, 21(6):860–868, 2018. doi:10.1038/s41593-018-0147-8. URL 10.1038/s41593-018-0147-8.

[26] Nathaniel D Daw, Samuel J Gershman, Ben Seymour, Peter Dayan, and Raymond J Dolan. Model-based influences on humans’ choices and striatal prediction errors. Neuron, 69(6):1204–1215, 2011.

[27] Silvia Maggi, Rebecca M Hock, Martin O’Neill, Mark Buckley, Paula M Moran, Tobias Bast, Musa Sami, and Mark D Humphries. Tracking subjects’ strategies in behavioural choice experiments at trial resolution. eLife, 13:e86491, mar 2024. ISSN 2050-084X. doi:10.7554/eLife.86491. URL 10.7554/eLife.86491.

[28] Jonathan D Cohen, Samuel M McClure, and Angela J Yu. Should I stay or should I go? How the human brain manages the trade-off between exploitation and exploration. Philosophical Transactions of the Royal Society B: Biological Sciences, 362(1481):933–942, March 2007. ISSN 0962-8436. doi:10.1098/rstb.2007.2098. URL 10.1098/rstb.2007.2098. _eprint: https://royalsocietypublishing.org/rstb/article-pdf/362/1481/933/93140/rstb.2007.2098.pdf.

[29] Kenji Doya, Kayoko W Miyazaki, and Katsuhiko Miyazaki. Serotonergic modulation of cognitive computations. Current Opinion in Behavioral Sciences, 38:116–123, 2021. ISSN 2352-1546. doi:10.1016/j.cobeha.2021.02.003. URL https://www.sciencedirect.com/science/article/pii/S2352154621000255. Computational cognitive neuroscience.

[30] Pierpaolo Iodice, Claudio Ferrante, Luigi Brunetti, Simona Cabib, Feliciano Protasi, Mark E Walton, and Giovanni Pezzulo. Fatigue modulates dopamine availability and promotes flexible choice reversals during decision making. Scientific Reports, 7(1):535, 2017.

[31] Patrick S. Hogan, Steven X. Chen, Wen Wen Teh, and Vikram S. Chib. Neural mechanisms underlying the effects of physical fatigue on effort-based choice. Nature Communications, 11(1):4026, 2020. doi:10.1038/s41467-020-17855-5. URL 10.1038/s41467-020-17855-5.

[32] Vivian V Valentin, Anthony Dickinson, and John P O’Doherty. Determining the neural substrates of goal-directed learning in the human brain. J Neurosci, 27(15):4019–4026, Apr 2007. ISSN 1529-2401 (Electronic); 0270-6474 (Print); 0270-6474 (Linking). doi:10.1523/JNEUROSCI.0564-07.2007.

[33] Wouter Kool, Joseph T McGuire, Zev B Rosen, and Matthew M Botvinick. Decision making and the avoidance of cognitive demand. J Exp Psychol Gen, 139(4):665–682, Nov 2010. ISSN 1939-2222 (Electronic); 0096-3445 (Print); 0022-1015 (Linking). doi:10.1037/a0020198.

[34] Amitai Shenhav, Matthew M. Botvinick, and Jonathan D. Cohen. The expected value of control: An integrative theory of anterior cingulate cortex function. Neuron, 79(2):217–240, 2013. ISSN 0896-6273. doi:10.1016/j.neuron.2013.07.007. URL https://www.sciencedirect.com/science/article/pii/S0896627313006077.

[35] Javier Masís, Travis Chapman, Juliana Y Rhee, David D Cox, and Andrew M Saxe. Strategically managing learning during perceptual decision making. eLife, 12:e64978, feb 2023.

[36] Marta Blanco-Pozo, Thomas Akam, and Mark E. Walton. Dopamine-independent effect of rewards on choices through hidden-state inference. Nature Neuroscience, 27(2):286–297, 2024. doi:10.1038/s41593-023-01542-x. URL 10.1038/s41593-023-01542-x.

[37] Peter Smittenaar, Thomas H. B. FitzGerald, Vincenzo Romei, Nicholas D. Wright, and Raymond J. Dolan. Disruption of dorsolateral prefrontal cortex decreases model-based in favor of model-free control in humans. Neuron, 80(4):914–919, 2025/12/06 2013. doi:10.1016/j.neuron.2013.08.009. URL 10.1016/j.neuron.2013.08.009.

[38] Wendy Wood and David T. Neal. A new look at habits and the habit-goal interface. Psychological review, 114 4: 843–63, 2007. URL https://api.semanticscholar.org/CorpusID:7468475.

[39] Wendy Wood and Dennis Rünger. Psychology of habit. Annual Review of Psychology, 67(Volume 67, 2016): 289–314, 2016. ISSN 1545-2085. doi:10.1146/annurev-psych-122414-033417. URL https://www.annualreviews.org/content/journals/10.1146/annurev-psych-122414-033417.

[40] Jianning Chen, Masakazu Taira, and Kenji Doya. Fictive learning in model-based reinforcement learning by generalized reward prediction errors. bioRxiv, 2025. doi:10.1101/2025.06.12.659433. URL https://www.biorxiv.org/content/early/2025/06/15/2025.06.12.659433.

[41] Sang Wan Lee, Shinsuke Shimojo, and John P O’doherty. Neural computations underlying arbitration between model-based and model-free learning. Neuron, 81(3):687–699, 2014.

[42] James G. March. Exploration and exploitation in organizational learning. Organization Science, 2(1):71–87, 1991. ISSN 10477039, 15265455. URL http://www.jstor.org/stable/2634940.

[43] K. Doya. Modulators of decision making. Nature Neuroscience, 11(4):410–6, 2008.

[44] Bruno B. Averbeck. Theory of choice in bandit, information sampling and foraging tasks. PLOS Computational Biology, 11(3):1–28, 2015. doi:10.1371/journal.pcbi.1004164. URL 10.1371/journal.pcbi.1004164.

[45] Tomohiko Yoshizawa, Makoto Ito, and Kenji Doya. Neuronal Representation of a Working Memory-Based Decision Strategy in the Motor and Prefrontal Cortico-Basal Ganglia Loops. eNeuro, 10(6):ENEURO.0413– 22.2023, 2023. doi:10.1523/ENEURO.0413-22.2023. URL http://www.eneuro.org/content/10/6/ENEURO.0413-22.2023.abstract.

[46] Peter S. Riefer, Rosie Prior, Nicholas Blair, Giles Pavey, and Bradley C. Love. Coherency-maximizing exploration in the supermarket. Nature Human Behaviour, 1(1):0017, 2017. doi:10.1038/s41562-016-0017. URL https://doi.org/10.1038/s41562-016-0017.

[47] Eric Schulz and Samuel J Gershman. The algorithmic architecture of exploration in the human brain. Current opinion in neurobiology, 55:7–14, 2019.

[48] Rahul Bhui, Lucy Lai, and Samuel J Gershman. Resource-rational decision making. Current Opinion in Behavioral Sciences, 41:15–21, 2021. ISSN 2352-1546. doi:10.1016/j.cobeha.2021.02.015. URL https://www.sciencedirect.com/science/article/pii/S2352154621000371. Value based decision-making.

[49] Wolfram Schultz, Peter Dayan, and P Read Montague. A neural substrate of prediction and reward. Science, 275 (5306):1593–1599, 1997.

[50] John P O’doherty. Reward representations and reward-related learning in the human brain: insights from neuroimaging. Current opinion in neurobiology, 14(6):769–776, 2004.

[51] Brian Knutson, Jonathan Taylor, Matthew Kaufman, Richard Peterson, and Gary Glover. Distributed neural representation of expected value. Journal of Neuroscience, 25(19):4806–4812, 2005.

[52] Kotaro Ishizu, Shosuke Nishimoto, Yutaro Ueoka, and Akihiro Funamizu. Localized and global representation of prior value, sensory evidence, and choice in male mouse cerebral cortex. Nature Communications, 15 (1):4071, 2024. ISSN 2041-1723. doi:10.1038/s41467-024-48338-6. URL 10.1038/s41467-024-48338-6.

[53] Ari Liu, Michael Schartner, International Brain Laboratory, and Ila Fiete. How learned expectations shape brain-wide responses. bioRxiv, 2025. doi:10.64898/2025.12.15.694430. URL https://www.biorxiv.org/content/early/2025/12/16/2025.12.15.694430.

[54] Roger Ratcliff. Group reaction time distributions and an analysis of distribution statistics. Psychological bulletin, 86 3:446–61, 1979. URL https://api.semanticscholar.org/CorpusID:13563153.

[55] Roger Ratcliff and Gail McKoon. The diffusion decision model: Theory and data for two-choice decision tasks. Neural computation, 20(4):873–922, 2008. ISSN 0899-7667 1530-888X. doi:10.1162/neco.2008.12-06-420.

[56] Roger Ratcliff, Philip L Smith, Scott D Brown, and Gail McKoon. Diffusion decision model: Current issues and history. Trends in cognitive sciences, 20(4):260–281, 2016.

[57] Milica Milosavljevic, Jonathan Malmaud, Alexander Huth, Christof Koch, and Antonio Rangel. The drift diffusion model can account for the accuracy and reaction time of value-based choices under high and low time pressure. Judgment and Decision making, 5(6):437–449, 2010.

[58] Satohiro Tajima, Jan Drugowitsch, and Alexandre Pouget. Optimal policy for value-based decision-making. Nature communications, 7(1):12400, 2016.

[59] Mads Lund Pedersen, Michael J Frank, and Guido Biele. The drift diffusion model as the choice rule in reinforcement learning. Psychonomic bulletin & review, 24(4):1234–1251, 2017.

[60] Laura Fontanesi, Sebastian Gluth, Mikhail S Spektor, and Jörg Rieskamp. A reinforcement learning diffusion decision model for value-based decisions. Psychonomic bulletin & review, 26(4):1099–1121, 2019.

[61] Nitzan Shahar, Tobias U Hauser, Michael Moutoussis, Rani Moran, Mehdi Keramati, Nspn Consortium, and Raymond J Dolan. Improving the reliability of model-based decision-making estimates in the two-stage decision task with reaction-times and drift-diffusion modeling. PLoS computational biology, 15(2):e1006803, 2019.

[62] Thomas Akam, Andy Lustig, James M Rowland, Sampath KT Kapanaiah, Joan Esteve-Agraz, Mariangela Panniello, Cristina Márquez, Michael M Kohl, Dennis Kätzel, Rui M Costa, et al. Open-source, python-based, hardware and software for controlling behavioural neuroscience experiments. Elife, 11:e67846, 2022.

[63] Michael K. Pitt and Neil Shephard. Filtering via simulation: Auxiliary particle filters. Journal of the American Statistical Association, 94(446):590–599, 1999. ISSN 01621459, 1537274X. URL http://www.jstor.org/stable/2670179.

[64] Randal Douc, Olivier Cappé, and Eric Moulines. Comparison of resampling schemes for particle filtering. CoRR, abs/cs/0507025, 2005. URL http://arxiv.org/abs/cs/0507025.

[65] Yoshihiko Ozaki, Shuhei Watanabe, and Toshihiko Yanase. OptunaHub: A platform for black-box optimization. arXiv preprint arXiv:2510.02798, 2025.

