## Supplemental material for "Continuous Strategy Adaptation and Discrete Switching Driven by Environment and Internal State in Meta-Learning"

### Supplementary material

#### 1 Generalized linear mixed model comparison

We validate the optimal structure by comparing several candidates of a generalized linear mixed model with a logit link function (GLMM) on simulated data, demonstrating the model's efficiency in differentiating between them (figure.1.b), and on experimental data, showing the model's fit to the data (figure.1.c). We included model-free strategy variants (MF, MF-memory, MF-loss, and MF-Prese), the model-based strategies (MB and LSI [1]), two hybrid model agents (MF-MB and MF-LSI), the MB-FL [2], and a random agent as the baseline (random) implemented as MF with near-zero inverse temperature.

Model 1 was originally designed to differentiate between model-based and model-free strategies by Out×Trans, which is positive after the trials when model-based strategies are more likely to stay (i.e., CR and RN), but model-free strategies are not. However, this term can yield false positives in pure model-free behavior, since choices leading to CR or RN trials are usually associated with a stronger reward history [1]. Model 2 improves the fit by introducing a "correct" term (i.e., whether the last choice is correct). Since the correct responses are often rewarded, it captures the reinforcing effect of past rewards, thereby isolating the genuine model-based consideration reflected by Out×Trans.

We further optimize Model 2. Firstly, the intercept term captures the baseline tendency toward choice repetition, yet the influence of choice perseveration is likely to increase with repeated presentations of identical content or over time. To account for possible accumulation of choice perseveration strength over time, we included the number of experienced trials as an additional regressor in Model 3, thereby improving model fit.

Moreover, "correct" information is unavailable to the animal, which reduces the psychological interpretability of the model. We replace it with the inferred value difference ($\Delta$Value), which is a cognitively plausible approximation of the effect of reward history. The variable is the difference in reward probabilities between the chosen and unchosen options estimated from the temporally discounted outcomes in history. Since it is derived from the outcome, we exclude it from the model to prevent collinearity. This regressor effectively substitutes for the function of "correct" and simplifies the model in Model 4 (figure.S1.b), resulting

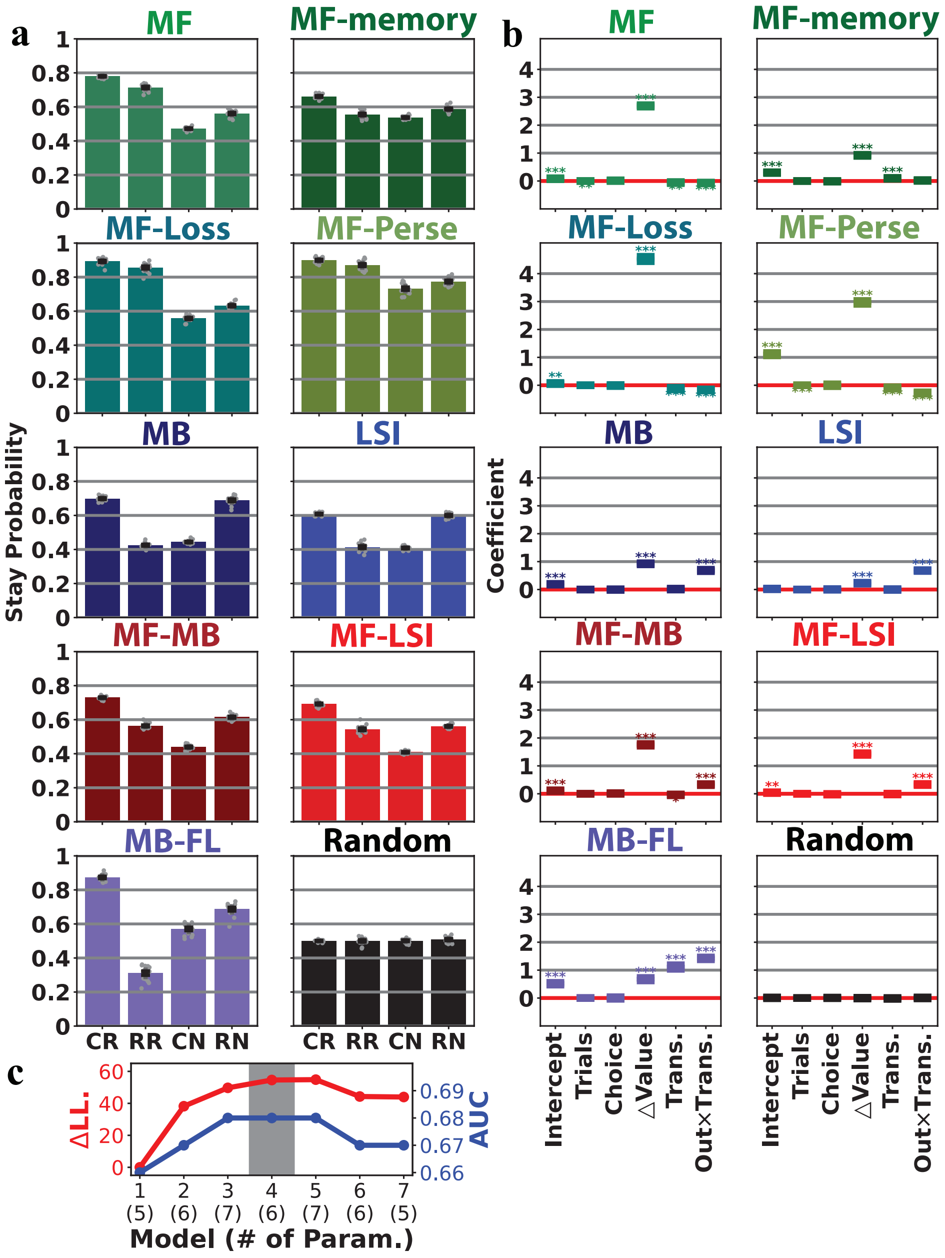


Figure 1: **a**, stay probability of simulation data each trial type (CR: common transition/reward; RR: rare transition/reward; CN: common transition/no-reward; RN: rare transition/no-reward).**b**. Results of GLMM model 4. Included coefficients are: intercept, the baseline of stay choice; Trials: the trial index; Choice, left (1) or right (0) choice; ΔValue: inferred value difference calculated as the difference in estimated reward probability of the previous chosen and unchosen option; Trans, transition type (rare: 0, common: 1); Out × Trans: the outcome transition type interaction (CR and RN: 1, CN and RR: 0). **c**, the model comparison by 10-fold cross-validation in real data. Δ LL. shows the difference in loglikelihood relative to model 1. AUC shows the area under the curve.

in a superior fit to actual behavior.

Including the entropy of the estimated reward probability of the chosen option marginally improves the fit at the expense of high model complexity in Model 5. Yet, GLMM shows no significant coefficient for it in either the simulation or the experiment data. Two extra models (Model 6 and 7) without the Trials regressor fit the real data worse, re-emphasizing the importance of the Trials term. Model 4 was selected because it can detect behavioral differences in strategies, fits the observed data well, and has a simple structure.

#### 2 Fixed-parameter modeling

We fit 61 RL models per session and subject separately and determine model fit using the Bayesian Information Criterion (BIC). For each session, $\Delta$ BIC was computed relative to the best-fitting model with the lowest BIC score in that session.

The included models are 11 base models and their variants with an additional learning rule. Based models include three model-free agents, two model-based agents, and their six hybrid models. Model-free learners update their expectations based on the mismatch between their expectations and actual outcomes, called prediction error; the extent of updating is determined by the learning rate $\alpha$. By having the eligibility trace between action and state $\lambda$, there are three model-free models: direct model-free learner without memory (MF), model-free learner with full eligibility trace between action and state (MF-memory), and a model-free learner with partial eligibility trace (MF-lambda). A model-based learner (MB) learns to map the action value to the state value by the action-state transition while learning the action value in a model-free manner. Besides, an agent might infer the latent state (which second-step state is good) in a Bayesian way, with a reversal probability $\psi$ [1]. Combining three model-free and two model-based learners gives six hybrid models, with $\epsilon$ governing the model-free contribution to the final decision relative to the model-based system.

Each of these base models is also combined with one of the five learning rules below. Loss aversion (Loss) treats the reward omission as a punishment with $\kappa$ scaling the strength of the punishment. Asymmetric learning learns the reward and omission event at different learning rates, $\alpha_{pos}$ and $\alpha_{neg}$. In parallel to learning, the forgetting rule (forget) states that the agent forgets the cached value for the choice that was not chosen or the state that was not experienced in the last trial, at the forgetting rate $\alpha_0$. Choice perseveration (Perse) maintains the choice kernel that tracks how often a choice has been chosen in the past by learning rate $\alpha_{CK}$, and its contribution to the final decision is weighted by $\epsilon_{CK}$. Fictive learning (FL) assumes that the agent estimate the correlation in reward contingency between experienced events (facts) and the events that were not experienced (counterfactual), $\eta$, and use this correlation to generalize the prediction error to update the counterfactual.

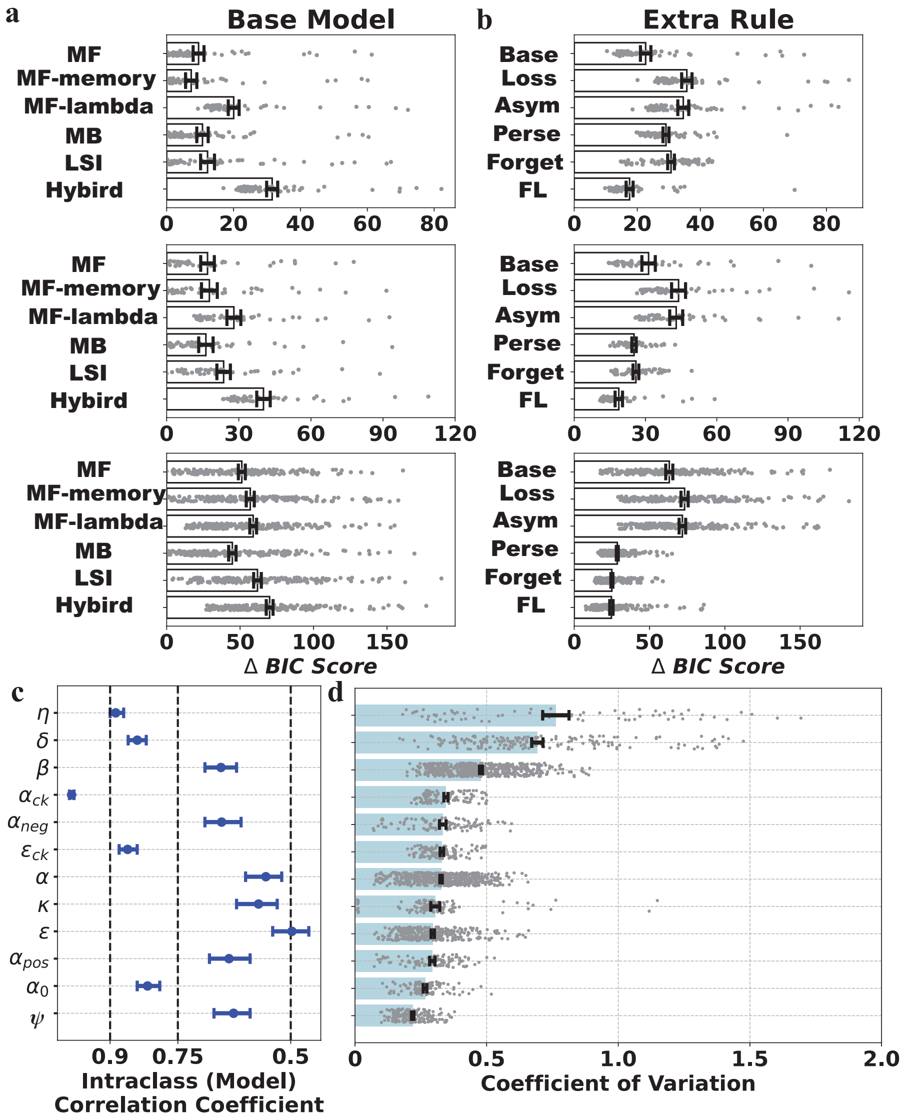


Figure 2: **Model fit diagnosis.a**, the Δ BIC score for model comparison in stage I (up), II (middle) and III (down) of the base models without additional rules. Included meta-parameters are: $\eta$ (event correlation for fictive learning), $\psi$ (environment change probability), $\beta$ (inverse temperature), $\alpha_{ck}$ (learning rate in choice perseveration), $\alpha_{neg}$ (learning rate for omission, only in asymmetric learning), $\epsilon_{ck}$ (weight of choice perseveration), $\alpha$ (learning rate), $\kappa$ (loss aversion strength), $\epsilon$ (weight of model-free relative to model-model system, model-free weight hereafter), $\alpha_{pos}$ (learning rate for reward, only in asymmetric learning), $\alpha_0$ (forgetting rate), and $\lambda$ (eligibility trace). **d**, the coefficient of variation of meta-parameters.

| parameter | $\mu_\rho$ | $\sigma_\rho$ | $p_{agreement}(\%)$ | $p_{significant}(\%)$ |
|---|---|---|---|---|
| $\beta$ | 0.7171 | 0.1460 | 100.0000 | 97.7049 |
| $\kappa$ | 0.3711 | 0.4053 | 85.4545 | 73.6364 |
| $\alpha_{\text{ck}}$ | 0.2157 | 0.2170 | 89.0909 | 20.0000 |
| $\lambda$ | 0.1472 | 0.2409 | 72.9412 | 18.2353 |
| $\alpha$ | 0.1018 | 0.3081 | 60.6897 | 26.0345 |
| $\alpha_{\text{neg}}$ | 0.0660 | 0.2546 | 56.0000 | 16.0000 |
| $\alpha_{\text{pos}}$ | -0.0395 | 0.2873 | 55.0000 | 25.0000 |
| $\alpha_0$ | -0.0406 | 0.2308 | 61.8182 | 9.0909 |
| $\epsilon_{\text{ck}}$ | -0.1443 | 0.3276 | 71.8182 | 40.0000 |
| $\epsilon$ | -0.2809 | 0.3584 | 79.6970 | 56.3636 |
| $\eta$ | -0.6453 | 0.1633 | 100.0000 | 97.1429 |
| $\psi$ | -0.7306 | 0.1210 | 100.0000 | 100.0000 |

Table 1: The summary of meta-parameter monotonicity by Spearman's correlation with session index. It displays the mean ($\mu_\rho$) and standard deviation ($\sigma_\rho$) of the correlation coefficient, along with the percentage of correlations in the same direction ($p_{agreement}(\%)$) and the percentage of significant coefficients ($p_{significant}(\%)$). The gray rows highlight the parameter that shows significant correlation in more than half of the sessions.

Consistent with the behavioral analysis, the model comparison revealed a transition over learning stages from model-free to model-based control, accompanied by the emergence of forgetting, choice perseveration, and fictive learning. MF(memory) best captured behavior in Stage I, but was overtaken by the classical model-based model, MB, in later stages (I: $w = 383, p < .0001$; II: $w = 487, p = 0.7373$; III: $w = 1918, p < .0001$, fig.2.a). Compared to the base models, the fictive learning rule improves the model fit consistently(I: $w = 132, p < .0001$; II: $w = 0, p < .0001$; III: $w = 0, p < .0001$), while forgetting (I: $w = 299, p < .0001$; II: $w = 374, p = 0.0.1070$; III: $w = 180, p < .0001$) and choice perseveration (I: $w = 492, p < .0001$; II: $w = 462, p = 0.5383$; III: $w = 401, p < .0001$) also improve the fit in stage III (fig.2.b).

The meta-parameter estimates are model-specific and notably vary across sessions (fig.2.c-d). We calculated the Intra-class Correlation Coefficient (ICC, two-way mixed effect, consistency type, single measurement [4]) across models to quantify the consistency of the meta-parameter estimates between models. ICC shows that 7 out of 12 meta-parameters have a poor to moderate level of agreement between models [3] (fig.2.c). The absolute Coefficient of Variation (CV) across sessions ($0.3854 \pm 0.2013$) shows significant variability (¿ 0.3) for most meta-parameters, suggesting noticeable change over sessions (fig.3.d). The Spearman's rank correlation coefficient analysis suggests that temporal change shows different levels of monotonicity across meta-parameters (table.1).

Hence, the model for dynamical investigation in the main text includes computational rules and meta-parameters that improve fit in fixed-parameter modeling. The model-based and model-free transitions suggest that the mixture model is indispensable. We include the eligibility trace (i.e., $\lambda$) to account for the mild transition from MF-memory to MF. Additionally, fictive learning (FL), forgetting (forget), and choice

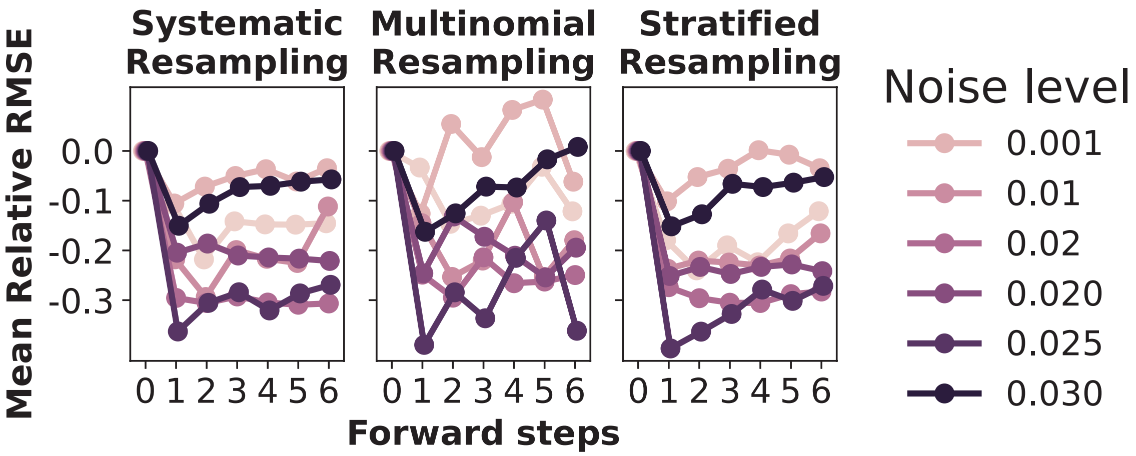


Figure 3: The improvement in root mean square error (RMSE) by including the forward step. The improvement is measured by the RMSE relative to it in the result without the forward step. The relative RMSE varies with the number of forward steps, the noise level, and the resampling method.

perseveration (Perse) are included.

#### 3 Validation of the forward steps in particle filtering

To validate whether the forward step we introduced improves the particle filtering performance. We simulate a hybrid model combining model-based and model-free RL with time-varying parameters and fit the generated data using particle filtering with different numbers of forward steps. The time-varying parameters are generated by a random walk with different levels of noise.

Results show that taking forward steps reduces the root mean square error (RMSE), but the optimal number of forward steps varies across noise level and resampling method (figure.3). The result validates that the forward step effectively improves the recoverability.

#### 4 Best-fitting hyperparameter in particle filtering

| hyperparameter | $\mu$ | $\sigma$ |
|---|---|---|
| $\alpha$ | 0.0213 | 0.0100 |
| $\beta$ | 0.0285 | 0.0154 |
| $\epsilon$ | 0.0276 | 0.0172 |
| $\lambda$ | 0.0275 | 0.0131 |
| $\eta$ | 0.0283 | 0.0138 |
| $\alpha_0$ | 0.0338 | 0.0149 |
| $\alpha_{ck}$ | 0.0197 | 0.0135 |
| $\epsilon_{ck}$ | 0.0251 | 0.0144 |

Table 2: The mean and standard deviations of the best-fitting hyperparameter in all subjects.

### 5 Behavioral characteristics across hidden states in FIS analysis

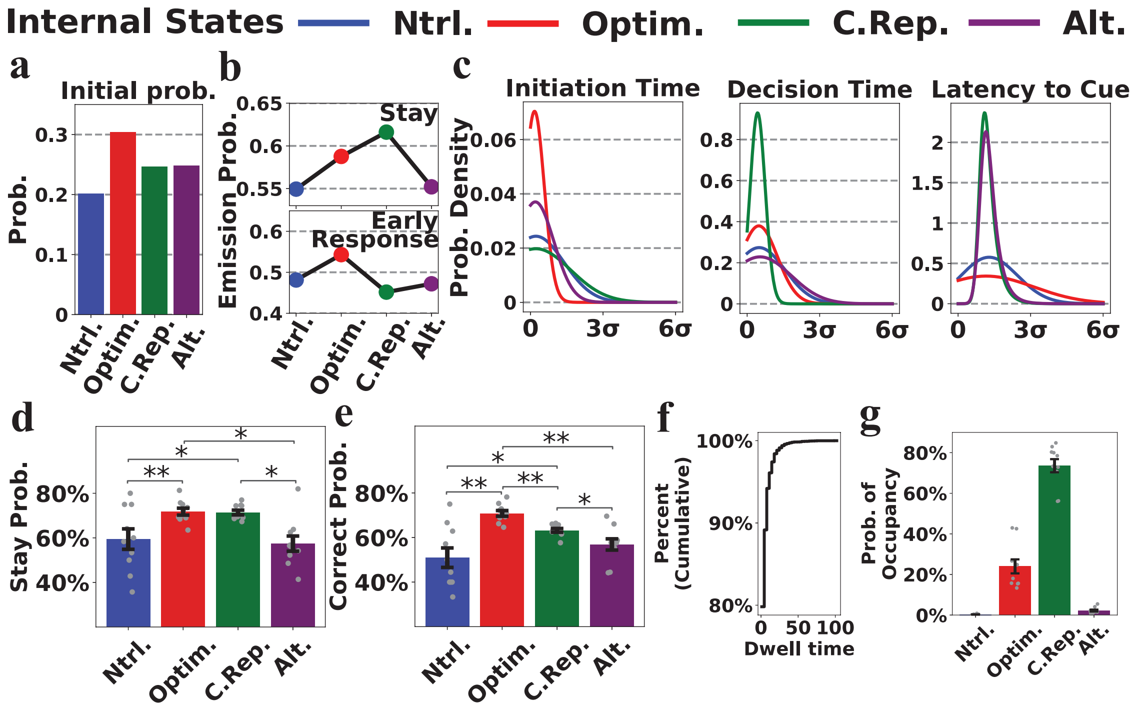


Figure 4: **The behavioral characteristic of four states.** Four states are neutral (Ntrl, blue), optimal (Optim., red), choice-repeating-impulsive (C.Rep., green), and alerted (Alt., purple) states. **a**, the initial probability of internal states. **b**, the estimated emission probability of stay choice (up panel) and early response (down panel). **c**, the probability density of initiation time, decision time, and latency to cue, reconstructed from the estimated exponential-Gaussian distribution parameter. **d, e**, the stay probability and accuracy from the recovered trajectory. **f**, the cumulative probability density of dwell time (i.e., number of trials before transition) in the recovered trajectory. **g**, the probability of occupancy in the recovered trajectory.

Here, we present an analysis of behavior across four states to better interpret the results.

The neutral state has the chance-level emission probability of stay (54.9389%, fig.4.b) and early response (48.2267%, fig.4.b), and the average-level distribution of the other three RT measures (fig.4.c). Animals achieved the lowest accuracy (50.9444% $\pm$ 13.8196%) here (fig.4.e). The alerted state is similar to the neutral state, except for the shorter latency to cue (fig.4.c), showing that an animal's response is primarily dominated by the reflex to the cue here. Because this distinction is limited to the state-check period, both stay probability and accuracy show no significant difference between the alerted and neutral states (fig.4.d&e), implying that both states represent different instances of disengagement.

The optimal state was characterized by the fastest initiation with the highest probability of early response, but exhibited prolonged decision time and latency to cue (fig.4.c), accompanied by superior performance (70.8143% $\pm$ 4.0460%) (fig.4.e). Animals promptly initiate the task to engage earlier, yet deliberate on the decision process and conserve energy by making slower cue responses, during which period a fast response has no extra gain. This state achieves an efficient trade-off between speed, accuracy, and cognitive load, representing optimal, accurate value-based learning.

The choice-repeating state exhibited the highest stay probability, accompanied by shorter decision time and latency to cue, but the slowest initiation time (fig.4.c), suggesting possible impulsiveness, awaiting further conclusive evidence. The high stay probability might be choice-repetition driven by choice perseveration, since the rapid decision prevents value-based deliberation, and this state shows lower accuracy ($63.2089\% \pm 2.6635\%$, fig.4.e). Since its mechanism remains unclear, we refer to this hidden state as the choice-repeating state, thereby distinguishing it from choice perseveration.

### References

[1] Thomas Akam, Rui Costa, and Peter Dayan. Simple plans or sophisticated habits? state, transition and learning interactions in the two-step task. *PLoS computational biology*, 11(12):e1004648, 2015.

[2] Jianning Chen, Masakazu Taira, and Kenji Doya. Fictive learning in model-based reinforcement learning by generalized reward prediction errors. *bioRxiv*, 2025. doi: 10.1101/2025.06.12.659433. URL `https://www.biorxiv.org/content/early/2025/06/15/2025.06.12.659433`.

[3] Terry K Koo and Mae Y Li. A guideline of selecting and reporting intraclass correlation coefficients for reliability research. *J Chiropr Med*, 15(2):155–163, Jun 2016. ISSN 1556-3707 (Print); 1556-3715 (Electronic); 1556-3707 (Linking). doi: 10.1016/j.jcm.2016.02.012.

[4] Kenneth O McGraw and Seok P Wong. Forming inferences about some intraclass correlation coefficients. *Psychological methods*, 1(1):30, 1996.
